# Targeted *in silico* characterization of fusion transcripts in tumor and normal tissues via FusionInspector

**DOI:** 10.1101/2021.08.02.454639

**Authors:** Brian J. Haas, Alexander Dobin, Mahmoud Ghandi, Anne Van Arsdale, Timothy Tickle, James T. Robinson, Riaz Gillani, Simon Kasif, Aviv Regev

## Abstract

Gene fusions play a key role as driver oncogenes in tumors, and their reliable discovery and detection are important for cancer research, diagnostics, prognostics and guiding personalized therapy. While discovering gene fusions from genome sequencing can be laborious and costly, the resulting “fusion transcripts” can be recovered from RNA-seq data of tumor and normal samples. However, alleged and putative fusion transcripts can also arise from multiple sources other than chromosomal rearrangements, including *cis*- or *trans*-splicing events, experimental artifacts during RNA-seq or computational errors of transcriptome reconstruction methods. Understanding how to discern, interpret, categorize, and verify predicted fusion transcripts is essential for consideration in clinical settings and prioritization for further research.

**Summary:** Here, we present FusionInspector for *in silico* characterization and interpretation of candidate fusion transcripts from RNA-seq and exploration of their sequence and expression characteristics. We applied FusionInspector to thousands of tumor and normal transcriptomes, and identified statistical and experimental features enriched among biologically impactful fusions. Through clustering and machine learning, we identified large collections of fusions potentially relevant to tumor and normal biological processes. We show that biologically relevant fusions are enriched for relatively high expression of the fusion transcript, imbalanced fusion allelic ratios, and canonical splicing patterns, and are deficient in sequence microhomologies detected between partner genes. We demonstrate that FusionInspector accurately validates fusion transcripts *in silico*, and helps identify and characterize numerous understudied fusions in tumor and normal tissues samples. FusionInspector is freely available as open source for screening, characterization, and visualization of candidate fusions via RNA-seq, and helps with transparent explanation and interpretation of machine learning predictions and their experimental sources.

**Highlights:**

- FusionInspector software for supervised analysis of candidate fusion transcripts
- Clustering of recurrent fusion transcripts resolves biologically relevant fusions
- Identification of distinguishing characteristics of known and novel fusion transcripts in tumor and normal tissues

## Introduction

Gene fusions are intensely studied for their relevance to disease and normal cellular biology. In cancer, gene fusions typically result from chromosomal rearrangements, including well-known drivers of cancer, such as BCR--ABL1 in chronic myelogenous leukemia (CML) (Kurzrock et al., 1988, Ren, 2005), TMPRSS2--ERG in prostate cancer (Tomlins et al., 2005, Rubin et al., 2011), and SS18--SSX1 or SS18--SSX2 in synovial sarcoma (Clark et al., 1994, Hale et al., 2019). Charting the diversity of fusion transcripts present in tumor and normal tissue is important for our basic understanding of the complexity and biological function of the transcriptome in normal and disease states, molecular diagnostics of cancer patients, and neoantigen discovery for targeting in personalized immunotherapy with cancer vaccines or T cell therapy (Yang et al., 2019, Wei et al., 2019).

The structural rearrangements leading to gene fusions can be detected or inferred through whole genome sequencing (WGS) or from the presence of “fusion transcripts” in whole transcriptome sequencing (RNA-seq) (Wang et al., 2013, Kumar et al., 2016a). Given the easier and economical nature of RNA-seq compared to WGS, and the effective methods for transcript assembly, RNA-seq has emerged as a leading experimental method for fusion transcript discovery and detection in both cancer research and molecular diagnostics. Dozens of computational tools have been developed to mine fusion transcripts from RNA-seq data (as referenced in (Haas et al., 2019)), and there have been multiple efforts to build catalogs of fusions across tumor and normal tissues (Klijn et al., 2015, Babiceanu et al., 2016, Hu et al., 2018, Yoshihara et al., 2015, Kim et al., 2010, Dehghannasiri et al., 2019). In general, tumor-specific fusion transcripts are presumed to derive from chromosomal rearrangements, whereas fusions identified in normal samples are considered more likely to be derived from normal karyotypes, thus reflecting other underlying causes, such as read-through transcription and *cis*- or *trans*-spliced products.

However, predicting fusions from RNA-seq data is challenging and the various methods developed to predict fusion products from RNA-seq vary tremendously in their accuracy for fusion detection, leading to both false positives and false negatives (Carrara et al., 2013, Kumar et al., 2016b, Haas et al., 2019). False positives can be driven by experimental artifacts that arise during reverse transcription or PCR amplification, and by computational mis-mapping of reads to target gene sequences (Yu et al., 2014), as well as specific differences in prediction tools. Moreover, as sequencing depth increases, the probability of detecting rare reads that support a fusion transcript prediction increases. This may be due to either lab artifacts or to real, low-rate trans-splicing of questionable functional relevance. Thus, there is an urgent need to understand the features that drive fusion detection and to generate high quality catalogs of well supported fusions.

Here, we describe FusionInspector (**Figure 1**), a method to assess and document the evidence for fusions. While other methods have previously been developed for supervised fusion detection and visualization (Schmidt et al., 2018, Schmidt et al., 2021, Lagstad et al., 2017, Kim et al., 2020, Zhang et al., 2017), FusionInspector includes specializations for comparing fusion transcripts to corresponding unfused fusion partners and aims to differentiate likely biologically relevant fusions from likely experimental or bioinformatic artifacts. FusionInspector reassesses the read alignment evidence supporting targeted candidate fusion transcripts, comparing the relative alignment evidence for a fusion transcript *vs*. counterevidence for its unfused partner transcripts. FusionInspector further evaluates the fusion transcript breakpoints in relation to sequence features representative of likely experimental and bioinformatic artifacts (Peng et al., 2015, Shivram and Iyer, 2018), including canonical splicing sequences, reference exon gene structures, and regions of microhomology between partner genes. Through reports, interactive visualizations, and classification, FusionInspector assists researchers in reasoning about the quantity and quality of the evidence supporting predicted fusions, to differentiate likely artifacts from fusions with characteristics similar to biologically relevant fusions known to occur in tumor and normal tissues.

**Figure 1.**
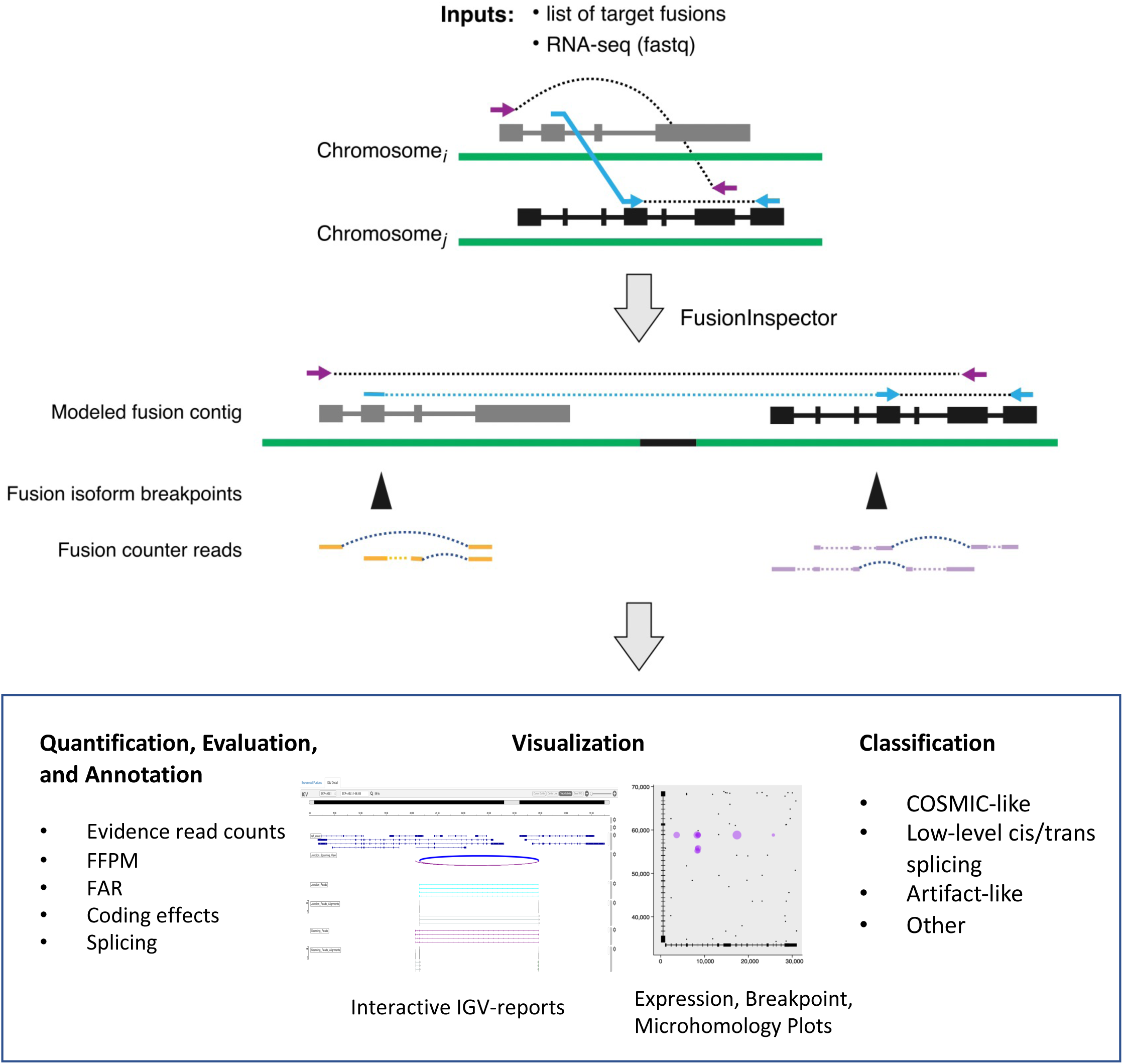
FusionInspector Overview. **Top:** Lists of fusion candidates derived from predictions of one or multiple fusion detection methods, or from a screening panel, are provided to FusionInspector as input along with RNA-seq in fastq format. For each candidate fusion gene, fusion contigs are generated by fusing the full-length gene candidates as collinear on a single contig. Intronic regions are by default each shrunk to 1 kb. RNA-seq reads are then aligned to a reference consisting of the entire genome supplemented with fusion contigs. Fusion-derived reads that would normally align discordantly as chimeric alignments in the reference genome (top example) instead align concordantly in the fusion contig context (bottom example). **Middle:** FusionInspector identifies split read alignments (light blue) and spanning pairs (purple) supporting the gene fusion in addition to read alignments that overlap the breakpoints and instead support the unfused fusion partners (fusion counter reads). **Bottom:** For those fusions where FusionInspector captures RNA-seq read support (“*in silico* validation”), it reports fusion sequence and expression characteristics, interactive visualizations, and predicted classifications as COSMIC-like, potential artifact, or other category (**Methods**).

We applied FusionInspector to assess recurrently predicted fusions in tumor and normal tissues, gathering insights into fusion transcript diversity, and devising machine learning methods for fusion classification (**Figure S1**). We first applied FusionInspector to examine recurrently predicted fusions in tumor and normal tissues to compute sequence and expression features, from which we next generated clusters of fusion isoforms with similar characteristics. We then identified fusion clusters that are enriched for known biologically relevant fusions, and others as likely artifacts. From these fusion clusters, we trained a classifier to automatically predict new fusion instances as likely biologically relevant, likely artifactual, derived from low-level cis- or trans-splicing, or other type. Finally, we applied FusionInspector on additional predicted instances of fusion transcripts that were clustered with biologically relevant fusions and explored, classified and prioritized these transcripts to identify new candidates in tumor and normal transcriptomes. Our application discovered a cluster of fusion transcripts heavily enriched for known cancer fusions and novel fusions of interest. FusionInspector is freely available as open source software at https://github.com/FusionInspector/FusionInspector/wiki.

## Results

### Development of FusionInspector for *in silico* evaluation of predicted fusion transcripts

FusionInspector (**Figure 1**) performs a supervised *in silico* evaluation of a specified set of candidate fusion transcripts, either predicted from RNA-seq data or from a user-defined panel. FusionInspector captures all read alignments in the RNA-seq that provide evidence for the specified fusions or for the unfused partner genes, and further explores the candidate fusion genes for regions of microhomology (defined as short identical sequence matches of length k (here *k*=10)), and the proximity of microhomologies to putative fusion breakpoints.

To capture evidence supporting candidate fusions, FusionInspector identifies those reads that align concordantly between fusion genes as juxtaposed in their fused orientation and provide concordant alignments that span the two genes in this rearranged context. There are two types of fusion-supporting alignments: (1) split-reads that define the fusion breakpoint, and (2) spanning fragments, where each paired-end read aligns to an opposite partner gene and the fragment bridges the fusion breakpoint (**Figure 1**). FusionInspector leverages STAR aligner (Dobin et al., 2013), which we enhanced here to support FusionInspector’s mode of action. As input to STAR, we provide the entire reference genome along with a set of fusion contigs constructed by FusionInspector (based on the list of specified fusion candidates), and STAR aligns reads to the combined genome targets and reports those aligned to the fusion contigs for further evaluation by FusionInspector (**Methods**).

Next, FusionInspector computes several features that are associated with the fusion based on these alignments and can assist in their evaluation. First, it uses the number of reads exclusively supporting each fusion as a proxy for the expression of the fusion transcript (similarly, read alignments overlapping the fusion breakpoint and exclusively supporting the unfused partner genes are a proxy for the expression levels of the unfused partner genes). Second, it computes the fusion allelic ratio (FAR) for the fusion with respect to each (5’ or 3’) partner transcript (5’-FAR and 3’-FAR) as the ratio of mutually exclusive reads supporting the fusion *vs*. each unfused partner gene (**Figures 1, S2)**. Third, it examines fusion breakpoints inferred from the read alignments for canonical dinucleotide splice sites at boundaries of the breakpoints in each partner gene and for agreement with available reference gene structure annotations. When there is evidence that supports multiple fusion transcript isoforms for a given fusion gene, FusionInspector uses an expectation maximization (EM)-based algorithm to fractionally assign mutually compatible spanning fragments to the corresponding isoforms (**Methods**). It then filters fusion candidates according to minimum evidence requirements (default settings require at least one split read to define the junction breakpoint, and at least 25 aligned bases supported by at least one read on both sides of the fusion breakpoint; **Methods**). Finally, it captures microhomologies between putative fusion genes and determines the proximity of a fusion breakpoint to the nearest site of microhomology.

By evaluating alignments to the modeled fusion contigs, FusionInspector demonstrates high sensitivity and specificity in supervised fusion detection, as we show for validated fusions in four well-studied breast cancer cell lines: BT474, KPL4, MCF7, and SKBR3. In our earlier study, 46 experimentally validated fusions were correctly predicted by at least one of 24 different fusion prediction methods (Haas et al., 2019). When FusionInspector was applied to the same data, it correctly identified each of these fusions in the relevant sample (**Figure 2, Supplementary Table S1**), with no false positives.

**Figure 2.**
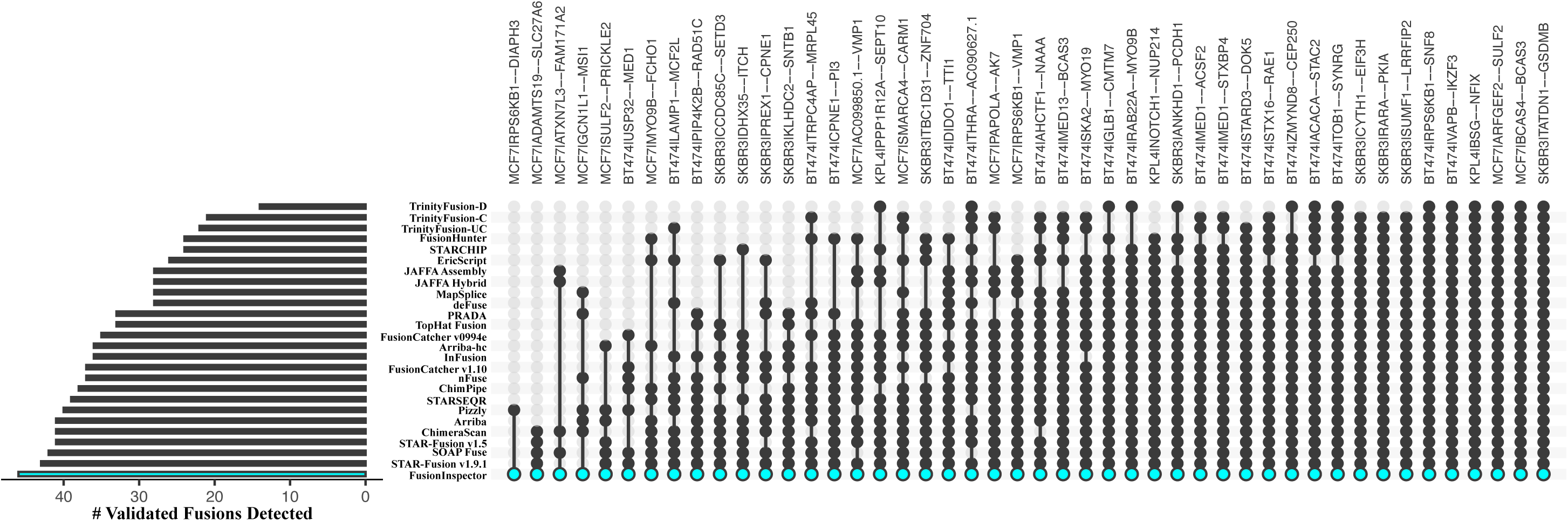
Detection of experimentally validated fusions in breast cancer cell lines BT474, MCF7, KPL4, and SKBR3. FusionInspector detects each of 46 experimentally validated fusions previously shown to be predicted by any of 24 different methods (Haas et al., 2019). FusionInspector results are highlighted as shown. While each sample was subject to inspecting identical lists of fusion candidates (Methods), fusions were specifically identified only in the cell lines for which they are known to exist.

Using sequence and expression attributes of fusions and known characteristics of biologically relevant fusions (below), FusionInspector further predicts whether each *in silico* validated fusion candidate is likely to be biologically relevant or alternatively has features consistent with experimental or bioinformatic fusion artifacts. We illustrate these features in the context of two contrasting examples of fusion types (**Figure 3**). Fusion EML4--ALK, a known cancer driver prevalent in lung adenocarcinoma (Soda et al., 2007, Sabir et al., 2017), has evidence of multiple transcript isoform structures, and while microhomologies are found between the EML4 and ALK genes, they tend to be distal from the fusion isoform breakpoints (**Figure 3a**). The EML4--ALK fusion breakpoints are all found at consensus dinucleotide splice sites that coincide with exon boundaries of reference gene structure annotations. In contrast, FusionInspector captures many reads supporting a putative fusion KRT13--KRT4, but the breakpoints inferred from split read alignments mostly have non-consensus dinucleotide splice sites and coincide with sites of microhomology, additionally, split reads with consensus dinucleotide splice sites mostly do not coincide with reference exon boundaries (**Figure 3b**). Because KRT13 and KRT4 are only distantly related with no easily detected nucleotide-level sequence conservation, their fusion may not be discarded by many fusion transcript predictors. However, given that most fusion evidence coincided with sites of microhomology and the lack of consensus splicing at breakpoints, FusionInspector infers most putative KRT13--KRT4 fusion isoforms to be artifactual. Another particularly compelling example of a similarly misleading and likely artifactual fusion is COL1A1--FN1, which is detected as prevalent in cancer-associated fibroblast cell lines (**Figure S3**). Further consideration of fusion and partner gene expression levels can aid in evaluating and prioritizing fusion candidates for further study, as we pursue below.

**Figure 3.**
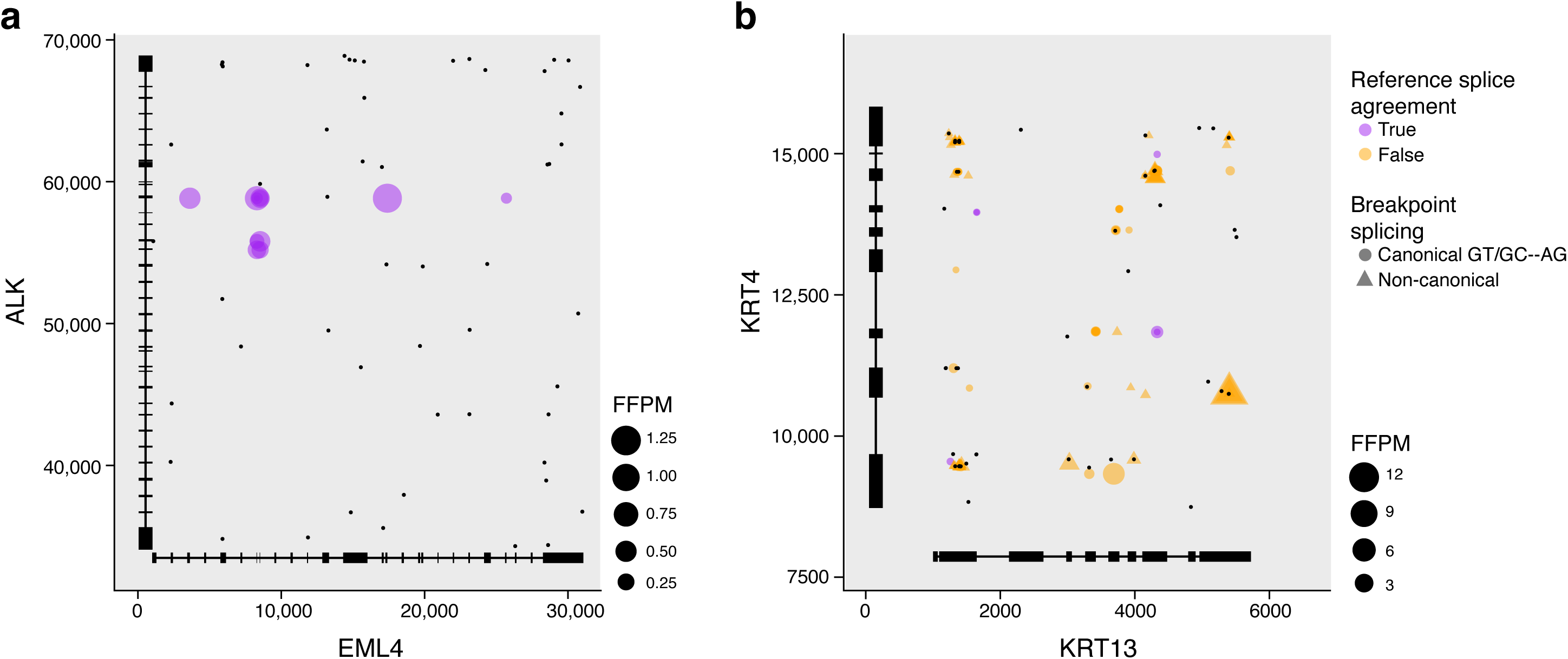
Features of fusion genes distinguish reliable and likely artifactual fusions. Fusion isoform expression level (dot size), splice type (dot color) and splice junction dinucleotide (dot shape) at each fusion breakpoint position involving the 5’ (x axis) and 3’ (y axis) partners of (**a**) EML--ALK (in COSMIC) and (**b**) KRT13--KRT4 (likely artifactual) fusions. Black dots: positions of microhomology (10 base exact match). Structures of collapsed isoforms for fusion partner genes are drawn along each axis.

### Clustering of recurrent fusion transcripts via FusionInspector attributes resolves COSMIC-like and artifact-like fusions

We first applied FusionInspector to examine recurrent fusion transcripts based on RNA-Seq from tumors (from TCGA (Cancer Genome Atlas Research et al., 2013)) and corresponding healthy tissue (from TCGA and GTEx (Consortium, 2013)). To generate an initial comprehensive catalog of input fusion isoforms in tumor and normal tissues, we predicted fusion transcripts with STAR-Fusion (v1.7) across 9,426 tumor and 707 normal samples from TCGA, and 8,375 normal samples from GTEx (**Table S2**). We initially applied lenient fusion evidence requirements to maximize sensitivity (**Methods**). As a result, putative fusion transcripts were detected in nearly all tumor and normal samples. After applying a minimum expression level threshold (0.1 FFPM), we detected a significantly higher number of fusions in tumors *vs*. paired normal samples in several TCGA tumor types (**Figure S4a**), although there were similar median numbers of predicted fusions per sample type in TCGA tumor and GTEx normal samples (t-test, p=0.5, **Figure S4b**). We readily identified known cancer fusions included in the COSMIC fusion collection (Forbes et al., 2017, “Wellcome Sanger Institute”, 2019) (“COSMIC-fusions”, **Figure 4a**) according to known disease associations and prevalence, such as TMPRSS2--ERG identified in roughly half of prostate cancers (Tomlins et al., 2005), FGFR3--TACC3 in glioblastoma (Lasorella et al., 2017), and PML--RARA in the acute promyelocytic leukemia subtype of acute myeloid leukemia (Liquori et al., 2020). COSMIC fusions were more highly expressed than most predicted fusions, which had low estimated expression levels and few supporting reads (**Figure 4b-d**).

**Figure 4:**
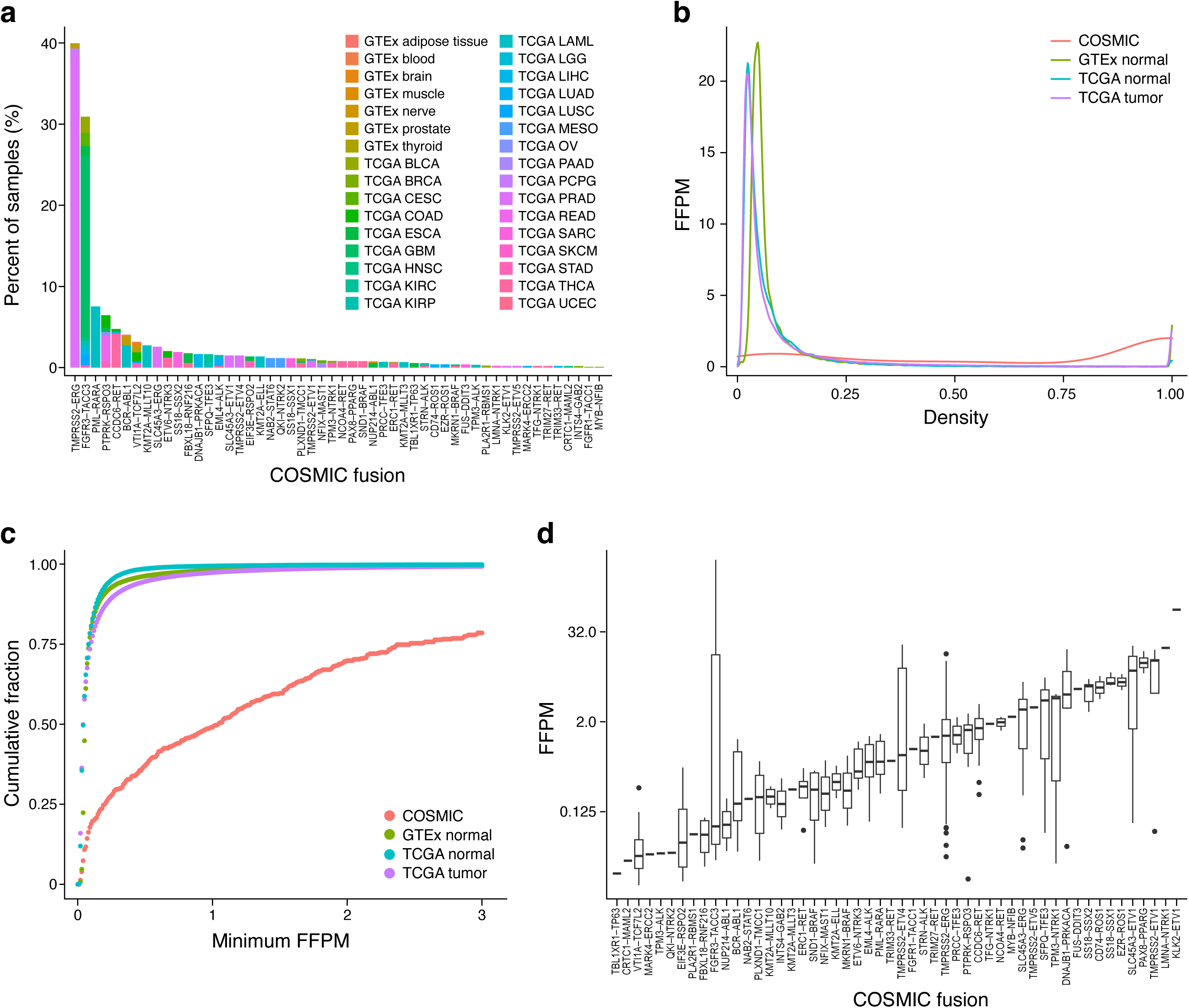
COSMIC fusions show distinctive properties among STAR-Fusion predictions across TCGA and GTEx. (**a**) Tissue and tumor composition. Percentages of TCGA tumor or GTEx normal samples (y axis) with corresponding predicted COSMIC fusions (x axis). TCGA study abbreviation codes as in (Commons”, 2021). (**b,c**) COSMIC fusions are more highly expressed than other predicted fusions. (**b-d**) Expression levels. (**b**) Distribution of fusion expression levels (*y* axis, FFPM; right-truncated at 1 FFPM) for all fusions predicted in TCGA tumors (purple), TCGA normal (blue), GTEx (green) and in COSMIC (red). (**c**) Cumulative fraction (*y* axis) of all predicted fusions at each minimum fusion expression (*x* axis, FFPM). (**d**) Distribution of fusion expression levels (y axis, FFPM) for each predicted COSMIC fusions (x axis). For a-d, fusions are restricted to the single highest expressed fusion isoform per sample occurrence, require reference annotation splice agreement at breakpoints, and have mitochondrial, HLA, and immunoglobulin gene containing fusions filtered.

Next, we used FusionInspector to examine the sequence and expression features of fusion transcripts that were recurrently detected across tumor and/or normal samples, in order to distinguish biologically impactful fusions (akin to the COSMIC fusions) from experimental or computational artifacts, or from low levels of *cis*- or *trans*-splicing from highly expressed genes. To this end, we analyzed 53,240 fusion isoforms (38,591 fusion occurrences and 14,649 alternatively spliced fusion isoforms) from 628 TCGA and 530 GTEx representative samples (**Methods**). For each fusion candidate, FusionInspector identified the number of reads supporting the fusion and those supporting the unfused partner genes at putative breakpoints, identified regions of microhomology between partner genes, and determined the following features: inferred fusion expression level (FFPM), 5’ and 3’ fusion allelic ratios (5’-FAR, 3’-FAR), 5’ and 3’ unfused gene expression levels (5’-counter-FFPM and 3’-counter-FFPM), presence of consensus *vs*. non-consensus dinucleotide splice sites at fusion breakpoints, agreement or disagreement with reference gene structure exon boundaries at splice junctions, number of microhomologies observed between the two partner genes, and the distance of each inferred fusion breakpoint to the nearest site of microhomology.

To distinguish fusion artifacts from those with features consistent with biologically impactful fusions, we clustered fusions by their feature profiles (**Figure 5a, Table S3, Methods**). Clustering produced 61 high granularity clusters, which we further grouped by hierarchical clustering according to median fusion attribute values in each fine cluster (**Figure 5b**). We then focused on examining clusters enriched for COSMIC fusions as a proxy for biologically impactful fusions.

**Figure 5.**
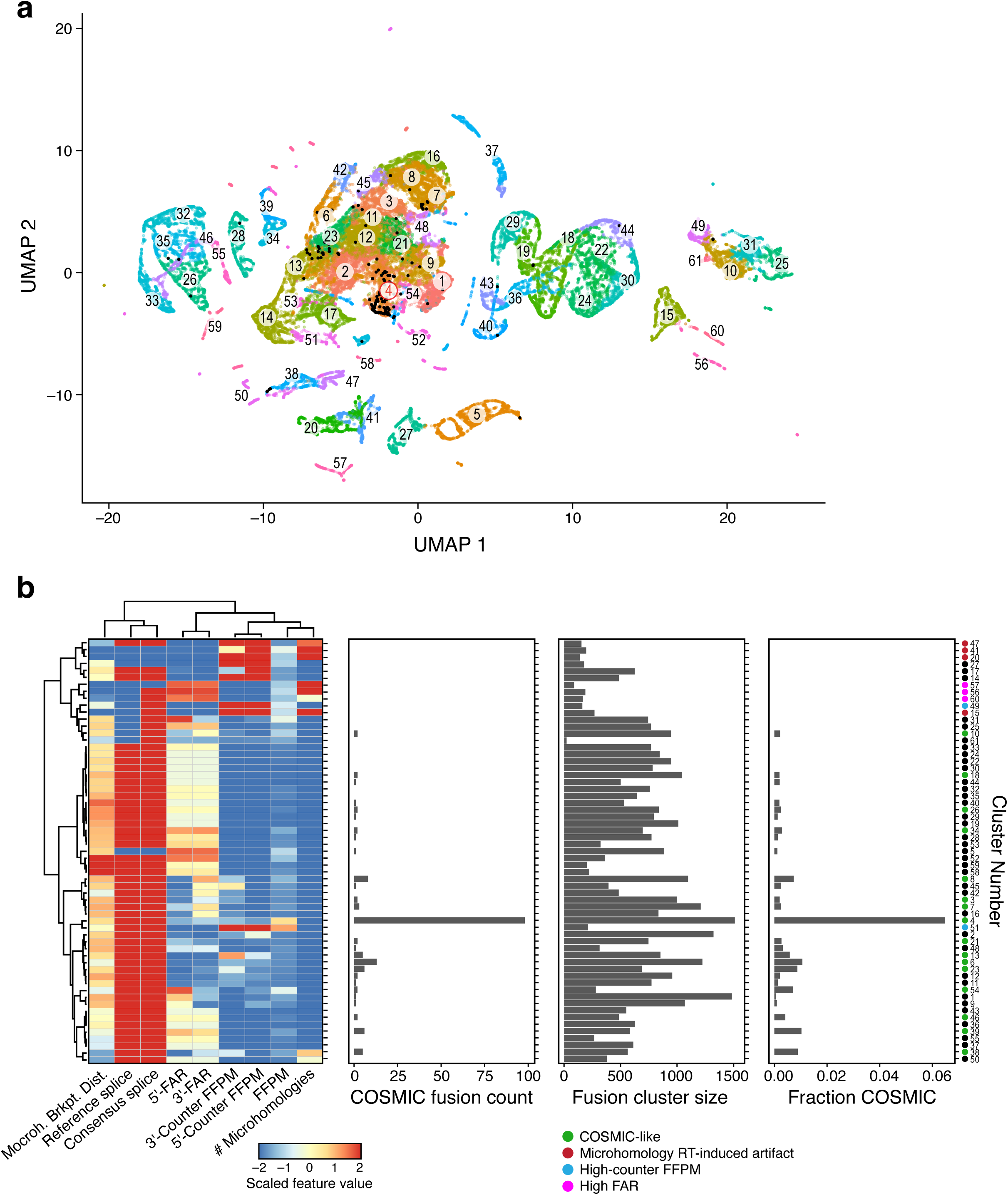
Fusions clustering by sequence and expression features distinguishes COSMIC-like fusions from likely artifactual ones. (**a**) Fusion clusters. Uniform Manifold Approximation and projection (UMAP) of 53,240 fusion isoforms feature profiles (dots), colored by Leiden cluster. Red label: Cluster C4. (**b**) Cluster C4 is enriched for COSMIC fusions. Features (columns, right), number of COSMIC fusions (x axis, second from left), cluster size (x axis, second from right), and fraction of COSMIC fusions (x axis, right) for each fusion cluster (rows). Heatmap shows median scaled intensity values for each feature (color bar).

One fusion cluster (C4) was significantly enriched with COSMIC fusions, harboring 57% of our detected instances of COSMIC fusions among these samples, but only 4% of all called fusions (p < 10^-90^, Fisher’s Exact one-sided test) (**Figure 5b**). Fusions in this cluster had splice breakpoints consistent with consensus splice sites and matching known reference gene structure exon boundaries, were relatively highly expressed, and were deficient in microhomologies between fusion partner genes. Most of the COSMIC fusions in C4 also have a 3’-FAR that exceeds the 5’-FAR, consistent with the fusion transcript being driven from an active 5’ partner’s promoter and a 3’ unfused partner expressed at lower levels (**Figure S5**). Sixteen additional clusters, all but one (C10) of which are members of one large hierarchical cluster with related features, had at least two COSMIC fusions per cluster and spanned 34% of the fusions overall and 37% of all COSMIC fusions.

Conversely, other fusion clusters, spanning 4% of all fusions and no COSMIC fusions (restricted to the highest expressed fusion isoform per occurrence), had features indicative of experimental or computational artifacts, especially enrichment in microhomologies that could confound alignment or contribute to RT mis-priming. We thus consider those fusions as putative artifacts. Of the 4% of all fusion occurrences encompassed by these clusters: half (2% of all fusions) had moderately-to-highly expressed partner genes, suggesting origination from RT mis-priming, and a 1/4 with little evidence for partner gene expression, suggesting read misalignments artifacts. The low portion of such presumed artifacts is a testament to STAR-Fusion’s rigorous filtering (Haas et al., 2019). The remaining 1/4 of putative artifact fusions involve highly expressed partner genes, where the detected fusion represented a small fraction of the total expression from these loci. These fusions may result from low levels of *cis*- or *trans*-splicing from the highly expressed partner genes.

### A fusion classifier allows targeted screening of predicted novel COSMIC-like fusions

We reasoned that the set of 1,511 predicted fusion occurrences (835 distinct gene pairings) that were members of the COSMIC-enriched cluster C4, are likely enriched for fusions of functional significance and should be prioritized for further study. Some are already known to be relevant to cancer but not yet included in the COSMIC database, such as EGFR--SEPT14 (Frattini et al., 2013), PVT1--MYC (Northcott et al., 2012, Jin et al., 2019, Tolomeo et al., 2021), and TPM3--NTRK1 (Ardini et al., 2014). Others are reciprocal fusions for COSMIC fusions that could result from balanced translocations, including reciprocal ABL1--BCR1 of COSMIC BCR1--ABL1, BRAF--SND1 of COSMIC SND1--BRAF, and PPARG--PAX8 of COSMIC PAX8--PPARG. This fusion cluster is also enriched for fusions exclusively identified in pancreatic tissue (explored below).

To gain further insights into the characteristics of the COSMIC-like fusions in C4, we first screened additional TCGA and GTEx samples to characterize additional occurrences of C4 representative fusions. We refocused FusionInspector on 236 C4+ key fusions (231 C4 recurrent fusion gene pairs including 26 COSMIC fusions, and another five COSMIC fusions not included in C4, **Methods, Table S4** (Haas, 2021)). We screened each of 2,764 TCGA and 1,009 GTEx representative samples for these 236 fusions (**Methods**), collecting FusionInspector validations and attributes for 37,211 additional fusion occurrences (**Figure 6**, **Figures S6a-e**, **Table S5**), and ranked fusions by the difference in their initial detected prevalence in tumors *vs*. normal samples.

**Figure 6.**
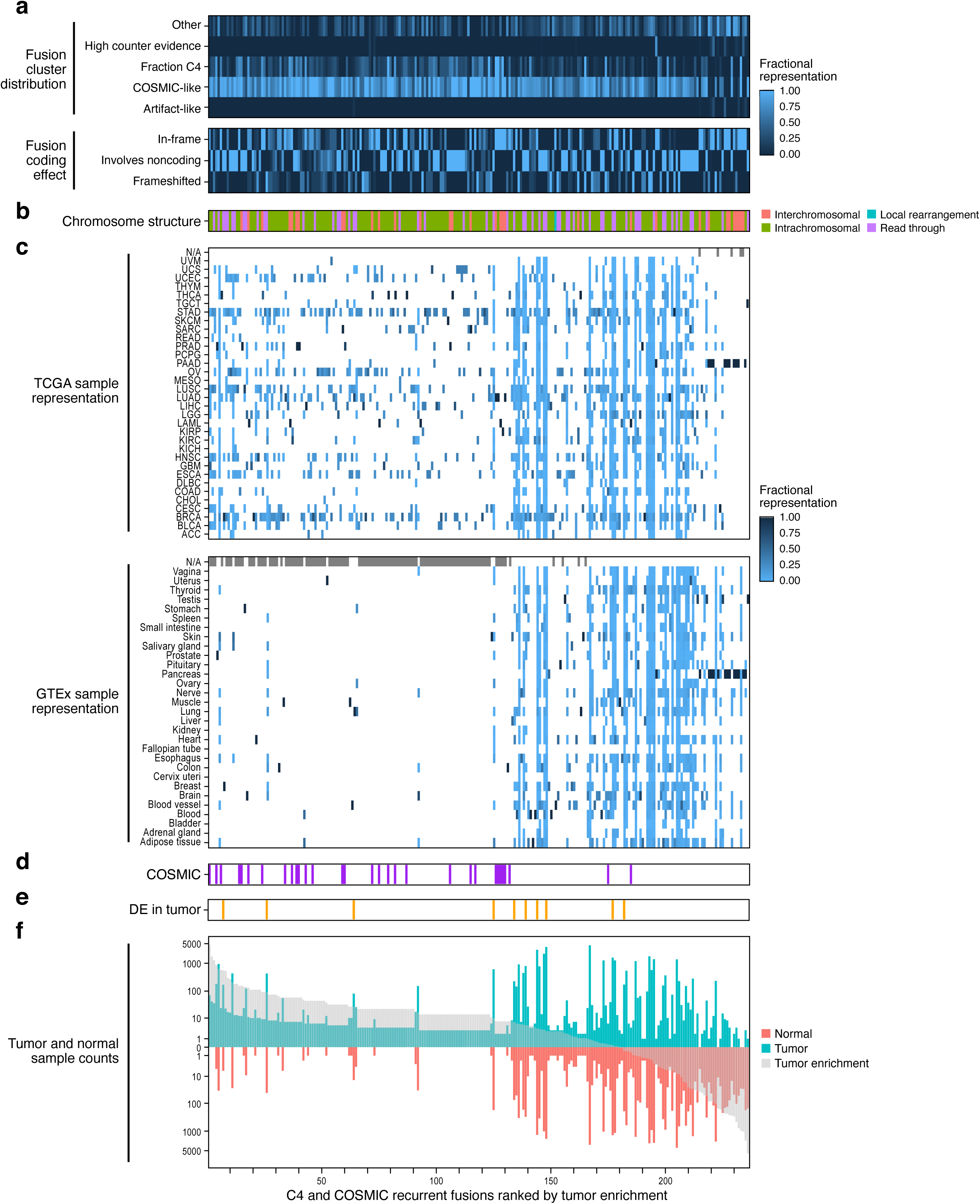
Characteristic properties of recurrent C4 and COSMIC fusions can distinguish biologically meaningful fusions and fusion instances. 236 selected COSMIC-peak-enriched (C4) and additional COSMIC fusions (columns / x axis) rank ordered by tumor enrichment and shown with fraction of the instances of each fusion in each category based on predicted Leiden cluster labels (a, rows, top) or corresponding to presumed impact on coding sequence (**a**, rows, bottom); fusion structure type based on the fusion partner’s chromosomal location (**b**); fraction of instances that is in each tumor or tissue type in TCGA and GTEx (**c**, rows); presence in COSMIC (**d**, purple), significantly higher expression in tumors *vs*. normal tissues (**e**, Wilcoxon rank sum test applied to FFPM, Benjamini Hochberg FDR < 0.05 and median tumor FFPM > median normal FFPM, orange), number of tumor (seagreen) or normal (light red) samples (f, y axis) predicted by STAR-Fusion to contain the fusion, rank ordered by tumor enrichment (f, x axis, (Methods, gray). Computations in **a**, **b**, and **d** are output by FusionInspector. See Figures S6a-e for each fusion pair.

To determine whether additional occurrences of these fusions have characteristics consistent with C4 fusions, general COSMIC-like fusions, are artifact-like, or belong to another category, we trained a random forest classifier to predict the labels of each of the earlier-defined 61 Leiden clusters and applied it to predict the cluster labels of fusion isoforms examined in this expanded targeted survey. We then categorized each fusion occurrence by the overall category (*e.g.*, “COSMIC-like”) of the hierarchical cluster to which the Leiden cluster label it was classified to belongs (**Figure 6a**, **Methods**). In this way, we distinguished individual fusion isoforms by their characteristics in the context of all the analyzed fusions. This random forest-based fusion classification was further incorporated into FusionInspector for routine application in discriminating COSMIC-like fusions from artifacts or other types.

FusionInspector-screened occurrences of known COSMIC fusions were mostly tumor-enriched with few to no normal samples identified with evidence (noting that all 1,009 GTEx normal samples were screened by FusionInspector for an identical list of COSMIC fusions). For most (27/31) known COSMIC fusions, at least 80% of fusion occurrences were classified as COSMIC-like, with the remaining four having at least half of occurrences classified as COSMIC-like, and none were classified as artifacts (**Figure 6a, Table 6S**). All but 31 of the 236 fusions had instances classified as C4, and all but 39 had at least 50% of their occurrences classified as COSMIC-like. Only 7 fusions had at least 10% of their occurrences classified as having high counter-evidence (C49 or C51), suggesting the fusion transcripts may reflect low levels of cis- or trans-splicing of more highly expressed normal fusion partner genes. Only 9 fusions had any occurrences predicted as artifacts, found in both TCGA and GTEx as pancreas-specific fusions (further discussed below).

This analysis highlighted intriguing, well-supported fusions for further study. For example, while the top-ranked tumor-enriched fusion, FGFR3--TACC3 (rank 1, 70 tumor samples, 0 normal) is a known oncogenic driver (Lasorella et al., 2017), other top ranking fusions, such as CCAT1-- CASC8 (rank 2, 42 tumors - mostly lung and stomach cancers, 0 normal) and VCL--ADK (rank 3, 36 tumors - also mostly lung and stomach cancers, 0 normal) have not yet been extensively studied. CCAT1--CASC8 was only recently reported in the fusion catalog generated by DEEPEST fusion (Dehghannasiri et al., 2019), and VCL--ADK was only previously reported in a study of cancer cell lines (Klijn et al., 2015).

### Some fusion transcripts are prevalent in normal tissues and may not be oncogenic

Approximately a third (86) of the 236 C4+ targeted fusions in our analysis were robustly detected in normal tissues (found in at least 5 normal samples); these may not be particularly relevant to cancer biology, but may play a role in normal biological processes. Of these, 61 fusions are broadly expressed across at least five tissues, involve intra-chromosomal pairs of genes, and can be largely explained by read-thru transcription, local rearrangements or *trans*- splicing of neighboring transcripts.

Some other putative fusions that are prevalent in normal tissues may in fact represent normal structural variation in the human genome, which is not accounted for when performing read alignment to a single human reference. For example, Fusion KANSL1--ARL17, which would require a local rearrangement in the human reference genome, is prevalent across both tumor and normal tissues (median of 31% of individuals, **Figure S7**), and is known to correspond to a common haplotype involving a locally rearranged genomic region observed in populations of European descent (Boettger et al., 2012). An earlier report identified KANSL1--ARL17 in diverse tumor samples and proposed that it may be a cancer predisposition germline fusion specific to Europeans (Zhou et al., 2017). Note, however, that no specific human genetic association evidence was shown for predisposition thus far, and we observe slightly higher prevalence of KANSL-ARL17 among GTEx normal samples than tumors from TCGA (**Figure S7**). Another normal fusion due to a rarer germline structural variation is TFG--GPR128, previously associated with a copy number variation and a haplotype frequency estimated at around 2% of individuals of European descent (Chase et al., 2010). Consistently, we find TFG- GPR128 broadly expressed across tumor and normal tissues and represented similarly at a median of 2% of all tissues examined (**Figure S8**). As more evidence of common structural variation becomes available, other prevalent fusions found in normal tissues may be more easily explained.

Another set of fusions that are less easily explained involve those we found only in normal pancreas and pancreatic carcinoma (**Figures 4, S6e**), involving various pairwise combination of CPA1, CPA2, CLPS, CELA2A, CELA3A, CTRB1, CTRB2, and CTRC (*e.g.*, CELA3A--CPA2, CELA3B--CELA2A, and CELA3A--CELA2A) fused to generate in-frame fusion products. These genes are among the highest expressed in pancreas and mostly on different chromosomes, suggesting trans-splicing may be the predominant underlying mechanism. While the transcripts were within the COSMIC-peak-enriched fusion cluster, the random forest based fusion classifier did not predict their newer instances as COSMIC-like, and some of them (*e.g.*, CELA3A-- CTRC) have high fractions of occurrences predicted as ’high-counter-evidence’ or ’artifact-like’ types (**Figures 6, S6e, Table 6S**).

### Some well-established oncogenic fusions are also reliably detected in normal samples

Several of the COSMIC fusions or other tumor-enriched fusions with known ties to cancer were surprisingly identified in both tumor and normal samples. For example, the prostate cancer fusion TMPRSS2—ERG, identified as our fourth most tumor-enriched fusion (182 of 465 TCGA prostate tumor samples), is also detected in six normal prostate samples (5 TCGA, 1 GTEx) (**Figure S6a**). TMPRSS2—ERG was only identified by FusionInspector in prostate tumors or normal prostate, reflecting both its high tumor-specificity and high specificity of fusion-calling.

In another example, COSMIC fusion PVT1--MYC was originally identified by STAR-Fusion in 21 samples (14 TCGA tumor, 1 TCGA normal, and 5 GTEx samples). Interactions between PVT1 and MYC including their fusion are well-known contributors to tumorigenesis (Northcott et al., 2012, Jin et al., 2019, Tolomeo et al., 2021). Through subsequent screening of PVT1-- MYC with FusionInspector, we identify a total of 32 samples (+9 TCGA tumor, +1 TCGA normal, and +3,-1 GTEx). Most (21/32) are expressed at low levels (below 0.1 FFPM), and we do not find strong evidence for expression to be generally higher in tumor samples than normals (p < 0.07, Wilcoxon rank sum test). However, five of the 32 PVT1--MYC occurrences were identified in cervical cancer tumors, and all were significantly more highly expressed than the other samples (p < 0.02), with the most highly expressed at 19 FFPM (**Figure S9**). PVT1 and MYC are co-localized to a proximal region in the bottom arm of chromosome 8 and a PVT1-- MYC fusion would likely involve local restructuring at the locus in tumors to generate the fusion product. Interestingly, this chromosome 8 region is a known hotspot for insertion of human papilloma virus (HPV) (Cancer Genome Atlas Research et al., 2017), the leading cause of cervical cancer (>90% of cases). Most of the TCGA cervical cancer samples we identified with PVT1--MYC have HPV insertions at this hotspot (see Table S3 of (Cancer Genome Atlas Research et al., 2017)). Thus, we hypothesize that HPV insertion contributes to the formation of the PVT1--MYC fusions. We did not find evidence for HPV insertion in the breast cancer sample with similar levels of PVT1--MYC expression (data not shown).

COSMIC fusion VTI1A--TCF7L2, originally identified as an oncogenic fusion in colorectal cancer (Bass et al., 2011), was most abundant in stomach, colon, and esophageal carcinoma samples, but also detected in seven individual GTEx normal samples (brain, whole blood, tibial nerve, tibial artery, prostate, and breast) (**Table S5**). While VTI1A--TCF7L2 was not enriched for detection in tumors *vs*. normal, only those fusions in colon cancer were highly expressed (>0.15 FFPM), whereas other tumor and normal instances were lowly expressed (<0.05 FFPM; many at the limit of detection; **Figure S10**), supported by a single split read defining the fusion breakpoint (**Table S5**). This could be consistent with a very low proportion of cells in the normal tissue expressing the fusion, compared to a large clone in the tumor.

Surprisingly, COSMIC fusion BCR--ABL1 was not tumor-enriched in our analysis, likely due to paucity of the relevant tumors in TCGA. In particular, BCR--ABL1 occurs in >95% of chronic myeloid leukemia (CML) cases (Kurzrock et al., 1988, Ren, 2005), but TCGA lacks CML samples. Indeed, the four TCGA tumors with BCR--ABL1 likely correspond to another subtype of AML defined with this fusion (Neuendorff et al., 2016). Three of these AML tumors also have evidence of the reciprocal ABL1--BCR fusion, and the oncogenic BCR--ABL1 is expressed at higher levels than the reciprocal counterpart in each sample. Interestingly, we detected eight instances of the oncogenic BCR--ABL1 fusion (and none of the reciprocal) in seven different GTEx normal tissues (1 each of adipose, breast, nerve, prostate, and thyroid and 2 pancreas). We observed no sequence or expression features distinguishing these fusions from those we identified in AML, and fusion breakpoints for those in GTEx normal tissues are identical to those found in AML. In general, when we find COSMIC fusions in GTEx normal samples, they are found at low frequencies (< 1% prevalence in a tissue type).

### Novel COSMIC-like fusion potentially relevant to breast and nasopharyngeal cancer

Other fusions clustered outside of C4 and found tumor enriched may also warrant further attention. For example, novel COSMIC-like fusion FSIP1--RP11-624L4.1 was detected in 22% of breast cancer tumors studied (240 of 1,086). While it was also detected in 14 normal breast samples (3 TCGA, 11 GTEx) and two additional samples (prostate and esophagus), its expression was significantly higher in tumors than normal tissue (**Figure 7a**, Benjamini Hochberg FDR < 0.004, Wilcoxon rank sum test). FSIP1 (fibrous sheath interacting protein 1) was previously identified as a prognostic marker for HER2-positive breast cancers and its high expression is associated with poor patient outcomes (Yan et al., 2019). Fusion partner RP11- 624L4.1 is a lncRNA, which is collinear and 170kb downstream from FSIP1, and was recently identified as an oncogene relevant to nasopharyngeal carcinoma (Zhou et al., 2020). FSIP1 and RP11-624L4.1 expression is positively correlated in both tumor and normal tissues (Pearson *r* = 0.6) and the fusion FSIP1--RP11-624L4.1 is found only among those samples most highly expressing both fusion partners (**Figure 7b**).

**Figure 7:**
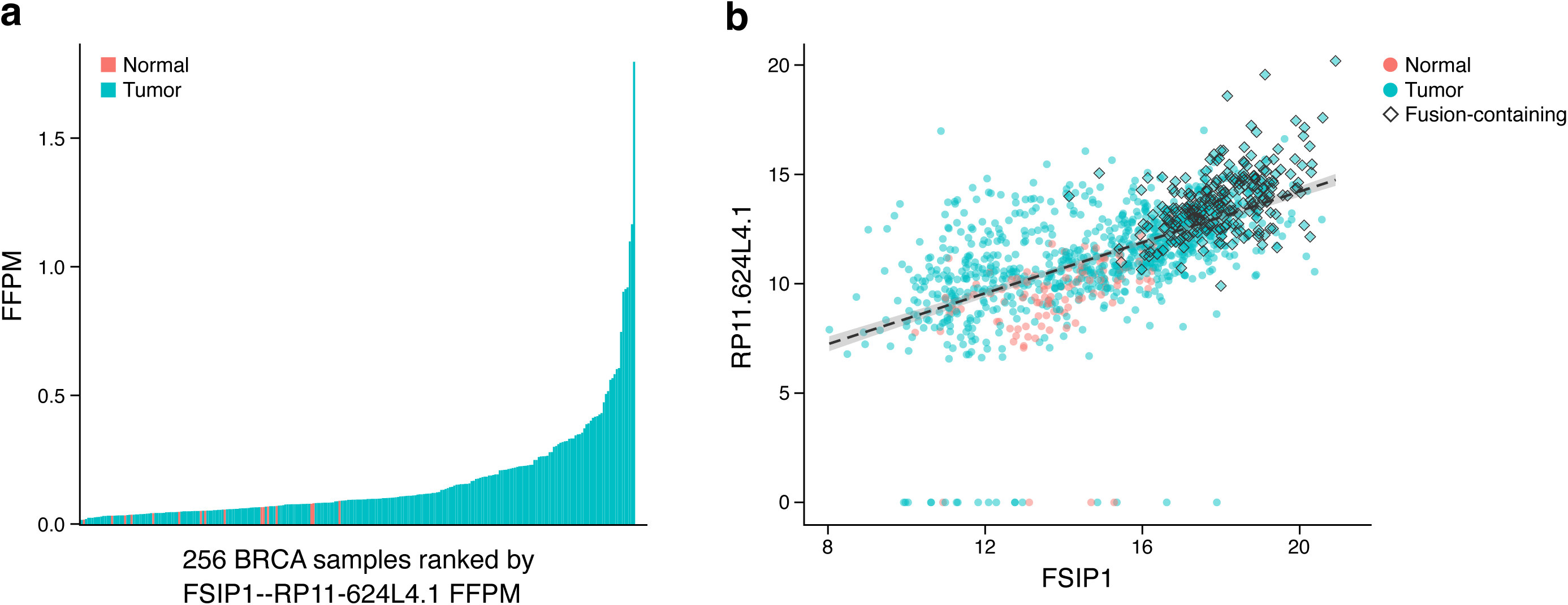
FSIP1--RP11-624L4.1 fusion in breast cancer. (**a**) Expression level (FFPM, y axis) of FSIP1--RP11-624L4.1 fusion in tumor (blue) and normal (red) TCGA breast cancer samples, ranked by FSIP1--RP11-624L4.1 expression. (**b**) Expression levels of FSIP1 (x axis) and RP11-624L4.1 (y axis) in each tumor (blue) and normal (red) samples (Pearson r=0.6, p-value < 2.2e- 16). Diamonds: samples where the fusion trsanscript is detected. Expression values were log_2_ transformed upper-quartile normalized gene FPKM measurements obtained from the Xena platform (Goldman et al., 2020).

## Discussion

We developed FusionInspector to enable exploration of the evidence supporting candidate fusions, flag likely artifacts, and identify those fusions with sequence and expression features similar to known biologically relevant fusion transcripts. Given a list of candidate fused gene pairs, FusionInspector captures RNA-seq read alignments that support either the fused genes or the unfused partner genes. From the fusion and partner gene expression evidence coupled with sequence features relating to the fusion breakpoint, FusionInspector helps the user to reason about the nature and quality of any target fusion transcript.

Clustering fusions by shared sequence and expression features identified a cluster of fusions highly enriched for COSMIC fusions. Fusions in this COSMIC “peak enriched” cluster had relatively high fusion expression with 3’-FAR generally exceeding 5’-FAR, suggesting oncogenic activity from the 3’-fused transcript. Analysis initiated by fusions in the COSMIC peak enriched cluster highlighted several putative novel or less appreciated oncogenic fusions, including CCAT1--CASC8 and VCL—ADK, based on their feature similarity to other well-known tumor-enriched fusions. Only ∼3% of initially predicted fusions were members of clusters likely enriched for artifacts based on features such as high partner gene expression or sites of microhomology at or near the fusion breakpoint. The low artifact rate is likely due to the strong filtering of the initial input catalog from STAR-Fusion.

While we focused on fusions identified in the COSMIC-peak-enriched cluster, other COSMIC-like fusion clusters also harbor important oncogenic fusions. For example, the COSMIC fusions SS18--SSX1 and SS18--SSX2, known drivers of synovial sarcoma (Clark et al., 1994, Gazendam et al., 2021), are in other clusters (C38 & C39), due in part to their higher 5’-FAR. Another fusion of interest, FSIP1--RP11-624L4.1 is present in 240 (22%) of breast tumors analyzed and in 16 normal breast tissues samples, where it is expressed at significantly lower levels. While the individual fusion partners have cancer associations (Yan et al., 2019, Zhou et al., 2020), any role for this newly identified fusion transcript deserves consideration in further exploring the roles of both genes in disease.

In some cases, fusions that were found in both tumor and normal tissues might reflect a low level of oncogenic events. For example, hallmark driver fusions including TMPRSS2--ERG and BCR--ABL1 are also detected in GTEx normal tissue samples, which may reflect low proportion of pre-malignant or transformed cells (Jaiswal and Ebert, 2019). We also detect COSMIC fusion VTI1A--TCF7L2 across multiple tumor and normal tissue types (consistent with (Nome et al., 2014)), but only highly expressed in colon cancer samples where it is a postulated oncogenic driver (Bass et al., 2011, Davidsen et al., 2018). Whether such a fusion could contribute to tumorigenesis in a different tissue with different cellular circuitry remains unknown.

Fusions that were prevalent among normal tissues can mostly be explained by read-through transcription and *cis*-splicing of colinear genes, but some may simply reflect natural germline structural variations that may exist in the population. With ongoing advancements in methods for detecting and cataloguing of structural variants (Collins et al., 2020, Abel et al., 2020), we may soon better understand the structural basis for many naturally occurring fusion transcripts. Access to matched RNA-seq and whole genome sequencing of the same samples across individuals would greatly facilitate such efforts.

Pancreas stood out as a clear outlier among all normal tissues explored for fusions. While we suspect some of the putative fusions detected in pancreas are derived from RT or alignment artifacts, several did have features consistent with *trans*-splicing of highly expressed partner genes, with *trans*-spliced products yielding in-frame proteins. In general, these fusion occurrences do not have COSMIC-like sequence and expression features. *Trans*-spliced in-frame fusion transcripts have the potential to expand functional diversity from our otherwise linear genomes (Gingeras, 2009), and even if these pancreas specific candidates failed to ultimately reach our COSMIC-like prioritization status, they may be worth additional studies.

FusionInspector opens the way to further explore the biological impact of the predicted fusions and the tissues and gene expression networks in which they are phenotypically relevant. FusionInspector helps illuminate the evidence supporting fusions in RNA-seq, or to sensitively and accurately screen for relevant fusions in samples of interest. Since short reads remain limited in their capacity to represent full length fusion transcripts, FusionInspector further integrates Trinity (Grabherr et al., 2011, Haas et al., 2013) for *de novo* reconstruction to optionally reconstruct more full-length fusion transcripts from RNA-seq data aligned to each fusion contig. FusionInspector is available as a stand-alone application for screening lists of candidate fusion transcripts, and is also incorporated into STAR-Fusion for *in silico* validation or visualization of STAR-Fusion predicted fusion transcripts. This facilitates analysis of fusions from both bulk and single cell RNA-Seq, as we have recently demonstrated (Jerby-Arnon et al., 2021).

Long read transcriptome sequencing may eventually obviate short read sequencing for fusion detection, thus removing the need for *de novo* reconstruction of full-length fusion transcripts (Liu et al., 2020, Rautiainen et al., 2020). Full-length single molecule direct RNA-Seq (Garalde et al., 2018) should also avoid RT amplification artifacts. Conversely, other features scored by FusionInspector, such as expression characteristics of fusion transcripts with respect to partner genes, will remain relevant and easily adapted for long read RNA-seq.

Machine learning is likely to play an increasingly important role in biomedical science and its clinical applications. In this paper we emphasize an important companion direction to machine learning, namely generating transparent and interpretable predictions, loosely referred to as explanations. The area of explanations and causal interpretation is growing rapidly in AI (Kasif and Roberts, 2020). We need to keep a reproducible trace of facts, predictions, and hypotheses from gene to function in the era of big data.

We hope that the practical applicability of FusionInspector will help drive transparency and other explanatory efforts in predictive areas in genomics and personalized medicine more generally, including screening and reassessment of evidence supporting fusion predictions, and visualization of the evidence via interactive reports (**Figures 1**, **S11**). FusionInspector can be easily leveraged as an add-on component to any fusion transcript prediction pipeline, and is directly incorporated into STAR-Fusion to facilitate execution as part of the Trinity Cancer Transcriptome Analysis Toolkit (Haas, 2019). The FusionInspector software is freely available as open source on GitHub, provided in container form via Docker and Singularity, and accessible on the Terra cloud computing framework for secure and scalable application across large compendiums of sample collections or patient-derived RNA-seq data.

## STAR Methods

### Key resources table

| Reagent or Resource | Source | Identifier |
| --- | --- | --- |
| **Deposited Data** |  |  |
| TCGA RNA-seq | http://Terra.bio | dbGaP: phs000854 |
| GTEX RNA-seq | http://Terra.bio | dbGaP: phs000424 |
| Breast cancer cell lines RNA-seq for BT474, KPL4, MCF7, and SKBR3 | (Edgren et al., 2011) | SRA:SRP003186 |
| **Software and Algorithms** |  |  |
| FusionInspector | This publication | https://github.com/FusionInspector/FusionInspector/wiki |
| STAR-Fusion | (Haas et al., 2019) | https://github.com/STAR-Fusion/STAR-Fusion/wiki |

### Resource availability

#### Lead contact

Further information and requests for resources or code should be directed to lead contact Brian Haas.

#### Materials availability

This study did not generate new unique reagents.

### Method Details

#### Initial comprehensive fusion transcript survey for TCGA and GTEx via STAR-Fusion

Fusions were predicted for TCGA and GTEx samples using STAR-Fusion (v1.7). First, the STAR (v2.6.1a) aligner was used to align RNA-seq reads from each sample to the human genome as follows:

”STAR --genomeDir ctat_genome_lib_build_dir/ref_genome.fa.star.idx --outReadsUnmapped None --chimSegmentMin 12 --chimJunctionOverhangMin 12 --chimOutJunctionFormat 1 -- alignSJDBoverhangMin 10 --alignMatesGapMax 100000 --alignIntronMax 100000 -- alignSJstitchMismatchNmax 5 -1 5 5 --runThreadN 16 --outSAMstrandField intronMotif -- outSAMunmapped Within --outSAMtype BAM Unsorted --readFilesIn reads_1.fastq reqds_2.fastq --outSAMattrRGline ID:GRPundef --chimMultimapScoreRange 10 -- chimMultimapNmax 10 --chimNonchimScoreDropMin 10 --peOverlapNbasesMin 12 -- peOverlapMMp 0.1 --genomeLoad NoSharedMemory --twopassMode Basic”.

The resulting Chimeric.out.junction files generated by STAR containing candidate chimeric reads were then analyzed by STAR-Fusion like so “STAR-Fusion -J Chimeric.out.junction –O $output_dir/STARF --genome_lib_dir $ctat_genome_lib --min_FFPM 0 --no_annotation_filter” leveraging CTAT genome library GRCh38_gencode_v22_CTAT_lib_Sept032019.

The parameters used here eliminated any filtering of fusions according to fusion expression levels or based on fusion annotations, so as to retain any fusions known to frequently occur in normal samples in both the normal and the tumor samples for further study. All STAR-Fusion predictions are provided in **Table S1**.

Fusion tumor enrichment was computed for each fusion according to ((# tumor with fusion + 1) / (total tumor samples)) / ( (# normal with fusion + 1) / (total normal samples) ).

#### FusionInspector method and implementation

FusionInspector takes as input a list of candidate fusions and RNA-seq files in fastq format, with each fusion formatted as “geneA--geneB” indicating a candidate fusion between geneA (5’) and geneB (3’). Leveraging the companion CTAT genome library set of genomic resources (identical to that used with STAR-Fusion, including the human reference genome, gene structure annotations, and STAR genome index), FusionInspector constructs fusion contigs by extracting the genomic sequences for each geneA and geneB, and concatenating each geneA and geneB pair into a single contig in collinear transcribed orientation. Gene structure annotations for fusion genes are similarly restructured to match the position and orientation of the corresponding genes in the fusion contigs. By default, long introns are shrunk to 1 kb in length by removing central regions of intron sequences, reducing the alignment search space and simplifying downstream visualizations.

RNA-seq reads are aligned to the fusion contigs along with the whole reference genome by running STAR with both inputs, including the pre-indexed whole genome and a fasta file containing the fusion contigs. STAR first loads the whole reference genome index into RAM, then builds an index for the fusion contigs, and incorporates the fusion contig index into the whole genome index. Only those reads that align concordantly to the fusion contigs, while considering all alignments to the combined targets, are reported. Note, that in the fusion context, all fusion-supporting reads are aligned concordantly, but will align partially to one gene and partially to the adjacent gene. This functionality was implemented in STAR since version 2.5.0a to support FusionInspector functionality. STAR-Fusion directly executes STAR to align reads like so “ STAR --runThreadN 4 --genomeDir ctat_genome_lib_build_dir/ref_genome.fa.star.idx --outSAMtype BAM SortedByCoordinate --twopassMode Basic --alignSJDBoverhangMin 10 -- genomeSuffixLengthMax 10000 --limitBAMsortRAM 47271261705 --alignInsertionFlush Right --alignMatesGapMax 100000 --alignIntronMax 100000 --readFilesIn reads_1.fastq.gz reads_2.fastq.gz --genomeFastaFiles finspector.fa --outSAMfilter KeepAllAddedReferences -- sjdbGTFfile finspector.gtf --alignSJstitchMismatchNmax 5 -1 5 5 --scoreGapNoncan -6 -- readFilesCommand ’gunzip -c’ “, where ’finspector.fa’ and ’finspector.gtf’ correspond to the fusion contigs sequence and structure annotation files.

FusionInspector examines the aligned reads output by STAR and identifies read alignments supporting fusions between gene pairs represented by the fusion contigs. Candidate fusion breakpoints are identified by split read alignments having partial alignments that anchor to exons of the neighboring fusion genes. Spanning fragments are identified as paired-end reads having each read mapping entirely on opposite sides of the breakpoint. Alignments must meet minimum evidence criteria to be counted as evidence, and require at least 96% sequence identity and no more than 10 bases unaligned at their ends (soft- or hard-clipped bases). For split reads, at least 10 bases must align adjacent to each breakpoint (anchor), and each anchor region must have sufficient sequence complexity, requiring entropy >= 1.2. For spanning fragments, each paired-end read must have sufficient sequence complexity, requiring entropy >= 1.2. Preliminary fusion predictions are defined based on candidate fusion breakpoints and sets of compatible spanning fragments. RNA-seq fragments that span a candidate breakpoint but support transcription from an unfused partner gene are captured and stored as counter-evidence and used to compute the partner gene counter FFPM and fusion allelic ratio.

There is often evidence for multiple fusion isoforms, and while the split reads are unique to and define each breakpoint, the spanning fragments are often compatible with multiple breakpoints and assigned ambiguously. We implemented an expectation maximization (EM) algorithm based on that described in kallisto (Bray et al., 2016) to fractionally assign RNA-seq evidence fragments to fusion isoforms according to maximum likelihood. Fusion expression values (FFPM) are then computed based on estimated RNA-seq fragment counts resulting from the EM. Fusion candidates are then filtered according to defined minimum evidence requirement, with defaults set as requiring at least one split read to define the junction breakpoint, and at least 25 aligned bases supported by at least one read on both sides of the fusion breakpoint. If the breakpoint involves non-consensus dinucleotide splice sites, then at least three split reads are required to support the breakpoint. Reads must also be found to align with at least 98% (default) sequence identity. A final filter of fusion predictions to exclude those containing overly promiscuous fusion partners (maximum 10) or those involving paralogs of more dominantly supported fusions is applied identically as previously described (Haas et al., 2017).

Optionally, Trinity de novo assembly (Grabherr et al., 2011, Haas et al., 2013) is integrated to de novo reconstruct candidate fusion transcripts based on reads aligning to the fusion contigs. When employed, Trinity-reconstructed fusion transcripts are identified in the final FusionInspector report and the assembled transcripts are available for further study. In addition, FusionInspector integrates IGV-reports (Robinson, 2019) to generate an interactive web-based summary (and fully self-contained html file) of predicted fusions coupled to a web-based interactive genome viewer to examine the read alignments found as evidence for the fusions.

#### Applications of FusionInspector to TCGA, GTEx, and cell lines

FusionInspector was run on TCGA v11 and GTEx v8 via Terra/AnVIL (Terra), as outlined in our analysis roadmap (**Figure S1**). First, FusionInspector v2.4.0 was used to reexamine a subset of 628 TCGA and 530 GTEx samples identified as containing instances of recurrent STAR-Fusion (v1.7) predictions. Candidate samples were identified based on individual fusions (a) having minimum 0.1 FFPM and (b) found in tissue types with at least three occurrences and comprising at least 10% of samples of that tissue type, or (c) containing a COSMIC fusion. Samples were then greedily selected to maximize recurrent fusion content while minimizing numbers of selected samples, retaining up to 10 samples per fusion. These samples were reexamined by executing the current STAR-Fusion (v1.9.1) including FusionInspector (v2.4.0) as a post-process like so: “STAR-Fusion --left_fq ${sample_name}_1.fastq --right_fq ${sample_name}_2.fastq --CPU 16 --genome_lib_dir ctat_genome_lib_build_dir --output_dir ${sample_name} --FusionInspector validate --no_annotation_filter --min_FFPM 0 “ leveraging companion CTAT genome library “GRCh38_gencode_v22_CTAT_lib_Apr032020”. The FusionInspector abridged outputs were consolidated and presented as **Table S3**. These fusions were subsequently subject to Leiden clustering (Traag et al., 2019) (see ***Fusion Clustering and Class Prediction*** section below).

Second, FusionInspector was run in fusion screening mode to explore instances of defined COSMIC-peak-enriched fusions (Leiden cluster 4 (C4) of the 61 fusion clusters found to be heavily enriched for COSMIC fusions). There were 231 instances of C4 fusions selected according to the following criteria: found in at least 3 samples, at least one fusion occurrence found clustered to C4, and at least 30% of occurrences annotated as COSMIC-like. These were further supplemented with five recurrent COSMIC fusions that are not members of C4 (ERC1-- RET, SLC34A2--ROS1, SS18--SSX1, SS18--SSX2, and VTI1A--TCF7L2), to a total of 236 fusion gene pairs (**Table S4**). The 236 fusion gene targets were provided as input to FusionInspector for screening 2,764 TCGA and 1,009 GTEx samples, each with the same list of 236 candidates. These samples were selected based on having a STAR-Fusion predicted occurrence of at least one of these fusions (from **Table S2**), and selecting a maximum of 50 samples per-fusion gene-pairing (with samples sometimes containing multiple fusion types), except for pancreatic and prostate cancer (TCGA) and normal pancreas tissue (GTEx) for which all samples were selected as targets. FusionInspector was executed like so: “FusionInspector -- fusions $Table_S4_fusions --genome_lib_dir ctat_genome_lib_build_dir -O ${sample_name} -- left_fq ${sample_name}_1.fastq --right_fq ${sample_name}_2.fastq --out_prefix ${sample_name} --vis” leveraging companion CTAT genome library “GRCh38_gencode_v22_CTAT_lib_Apr032020”, and results for screening of these samples are provided in **Table S5**.

Using RNA-seq data for breast cancer cell lines BT474, MCF7, KPL4, and SKBR3 as in (Haas et al., 2019), FusionInspector was run on each sample with a targeted list of 52 experimentally validated fusions (**Table S7**). Results for each sample are provided in **Table S1**.

#### Fusion transcript clustering and attribute class prediction

All 53,240 fusion isoforms surveyed by FusionInspector from our initial subset of TCGA and GTEx samples were clustered according to sequence and expression characteristics. Microhomologies defined as exact *k*-mers with *k*=10 were identified between candidate fusion gene pairs as represented in the FusionInspector-constructed fusion contigs (with introns shrunk to a max of 1 kb each for simpler visualizations). The Euclidean distance of each candidate fusion breakpoint to the nearest site of microhomology was determined in the FusionInspector fusion contig coordinate system. Attributes of interest for clustering fusions were: (1) the fusion expression level (FFPM), (2,3) partner gene fusion allelic ratios (5’-FAR and 3’-FAR), (4,5) the left and right unfused partner gene expression levels expressed as 5’- and 3’-counter-FFPM and computed based on the number of counter-reads observed as aligned at each corresponding gene breakpoint site, (6,7) indicators for consensus dinucleotides and agreement with reference gene structure exon boundaries at the fusion breakpoints, and (8) the number of microhomologies and (9) distance of the breakpoint to the nearest microhomology. These numerical values were centered and scaled to Z-scores, truncated within the interval [-2,2] to remove outliers, and then rescaled so each attribute numerical vector would fill the interval [-2,2] simplifying our evaluation of metrics using a consistent low-to-high range for each attribute type.

We calculated the distance between fusions based on vectors with these values, constructed a *k*- nearest-neighbor graph (*k*=50) of fusions, and clustered the graph by Leiden clustering (Traag et al., 2019) (resolution_parameter = 3). The impact of the resolution parameter on clustering and COSMIC fusion enrichment was examined (**Figure S12**), and the parameter with the most granular set of clusters was selected for further analysis. Clusters were manually reviewed and grouped and annotated according to median cluster attributes, with cluster annotation term assignments as “COSMIC-like” if clusters contained at least two COSMIC fusions, “COSMIC- peak-enriched” if predicted as cluster C4, “high-counter FFPM” indicating relatively high expression of the partner genes and potentially resulting from a low rate of trans-splicing, and categories “High FAR” and “Microhomology RT-induced artifact” to reflect likely bioinformatic or reverse-transcription related artifacts (as labeled in **Figure 5b**).

A random forest classifier was built to predict Leiden cluster membership based on scaled fusion attributes. The classifier was constructed by randomly selecting a maximum of 300 fusions (median cluster size) from each cluster, and leveraging 2/3 of fusions for training and 1/3 for testing, all performed using Ranger (Wright and Ziegler, 2017). Fusions predicted to be assigned to any cluster noted earlier with a fusion cluster annotation (*e.g.*, “COSMIC-like”) are assigned a prediction according to that fusion cluster annotation term. Such fusion attribute cluster predictions are now incorporated into the latest FusionInspector (v2.6.0).

#### Data and code availability

- FusionInspector source code is available on GitHub at: https://github.com/FusionInspector/FusionInspector
- Data are provided in supplementary tables in addition to being available on GitHub with code to demonstrate analysis methods and generation of figures (Haas, 2021) .
- RNA-seq fastq files for breast cancer cell lines BT474, MCF7, KPL4, and SKBR3 as leveraged here and in (Haas et al., 2019) are available at https://data.broadinstitute.org/Trinity/CTAT_FUSIONTRANS_BENCHMARKING/on_cancer_cell_lines/reads/

## Acknowledgements

We thank Joshua Gould for assisting with software pipeline development and Terra integration, and Leslie Gaffney for help with generating publication quality figures. This work has been supported by National Cancer Institute grants U24CA180922 (A.R. and B.J.H.), R50CA211461 (B.J.H.), Howard Hughes Medical Institute (A.R.), Klarman Cell Observatory (A.R.), NHGRI R01HG009318 (AD), and American Association of Obstetricians and Gynecologists Foundation (AAOGF) Scholar Grant 2017-2020 (AVA). The results published here are in whole or part based upon data generated by the TCGA Research Network: https://www.cancer.gov/tcga. TCGA v11 data were obtained through Terra/Anvil via dbGaP accesssion number phs000178.v11.p8. The Genotype-Tissue Expression (GTEx) Project was supported by the Common Fund of the Office of the Director of the National Institutes of Health, and by NCI, NHGRI, NHLBI, NIDA, NIMH, and NINDS. The GTEx v8 data used for the analyses described in this manuscript were obtained from Terra/AnVIL via via dbGaP accession number phs000424.v8.p2.

## Author Contributions

BJH performed analyses, developed the FusionInspector software, and wrote the initial draft of this manuscript. AD enhanced the STAR aligner software to support FusionInspector execution. JTR and TT developed the FusionInspector IGV-reports interactive fusion evidence visualization component. AVA assisted with investigations of HPV insertions and MYC--PVT1 fusion studies in CESC samples. AR and SK advised this work, and all authors made intellectual contributions towards the design of FusionInspector and to the final manuscript.

## References

1. ”WELLCOME SANGER INSTITUTE”. 2019. COSMIC Catalogue of Somatic Mutations in Cancer [Online]. Available: https://cancer.sanger.ac.uk/cosmic [Accessed].

2. Abel, H. J., Larson, D. E., Regier, A. A., Chiang, C., Das, I., Kanchi, K. L., Layer, R. M., Neale, B. M., Salerno, W. J., Reeves, C., Buyske, S., Genomics, N. C. F. C. D., Matise, T. C., Muzny, D. M., Zody, M. C., Lander, E. S., Dutcher, S. K., Stitziel, N. O. & Hall, I. M. 2020. Mapping and characterization of structural variation in 17,795 human genomes. Nature, 583, 83–89.

3. Ardini, E., Bosotti, R., Borgia, A. L., DE Ponti, C., Somaschini, A., Cammarota, R., Amboldi, N., Raddrizzani, L., Milani, A., Magnaghi, P., Ballinari, D., Casero, D., Gasparri, F., Banfi, P., Avanzi, N., Saccardo, M. B., Alzani, R., Bandiera, T., Felder, E., Donati, D., Pesenti, E., Sartore-Bianchi, A., Gambacorta, M., Pierotti, M. A., Siena, S., Veronese, S., Galvani, A. & Isacchi, A. 2014. The TPM3-NTRK1 rearrangement is a recurring event in colorectal carcinoma and is associated with tumor sensitivity to TRKA kinase inhibition. Mol Oncol, 8, 1495–507.

4. Babiceanu, M., Qin, F., Xie, Z., Jia, Y., Lopez, K., Janus, N., Facemire, L., Kumar, S., Pang, Y., Qi, Y., Lazar, I. M. & Li, H. 2016. Recurrent chimeric fusion RNAs in non-cancer tissues and cells. Nucleic Acids Res, 44, 2859–72.

5. Bass, A. J., Lawrence, M. S., Brace, L. E., Ramos, A. H., Drier, Y., Cibulskis, K., Sougnez, C., Voet, D., Saksena, G., Sivachenko, A., Jing, R., Parkin, M., Pugh, T., Verhaak, R. G., Stransky, N., Boutin, A. T., Barretina, J., Solit, D. B., Vakiani, E., Shao, W., Mishina, Y., Warmuth, M., Jimenez, J., Chiang, D. Y., Signoretti, S., Kaelin, W. G., JR., Spardy, N., Hahn, W. C., Hoshida, Y., Ogino, S., Depinho, R. A., Chin, L., Garraway, L. A., Fuchs, C. S., Baselga, J., Tabernero, J., Gabriel, S., Lander, E. S., Getz, G. & Meyerson, M. 2011. Genomic sequencing of colorectal adenocarcinomas identifies a recurrent VTI1A-TCF7L2 fusion. Nat Genet, 43, 964–968.

6. Boettger, L. M., Handsaker, R. E., Zody, M. C. & Mccarroll, S. A. 2012. Structural haplotypes and recent evolution of the human 17q21.31 region. Nat Genet, 44, 881–5.

7. Bray, N. L., Pimentel, H., Melsted, P. & Pachter, L. 2016. Near-optimal probabilistic RNA-seq quantification. Nat Biotechnol, 34, 525–7.

8. CANCER GENOME ATLAS RESEARCH, N., ALBERT EINSTEIN COLLEGE OF, M., ANALYTICAL BIOLOGICAL, S., BARRETOS CANCER, H., BAYLOR COLLEGE OF, M., BECKMAN RESEARCH INSTITUTE OF CITY OF, H., BUCK INSTITUTE FOR RESEARCH ON, A., CANADA’S MICHAEL SMITH GENOME SCIENCES, C., HARVARD MEDICAL, S., HELEN, F. G. C. C., RESEARCH INSTITUTE AT CHRISTIANA CARE HEALTH, S., HUDSONALPHA INSTITUTE FOR, B., ILSBIO, L. L. C., INDIANA UNIVERSITY SCHOOL OF, M., INSTITUTE OF HUMAN, V., INSTITUTE FOR SYSTEMS, B., INTERNATIONAL GENOMICS, C., LEIDOS, B., MASSACHUSETTS GENERAL, H., MCDONNELL GENOME INSTITUTE AT WASHINGTON, U., MEDICAL COLLEGE OF, W., MEDICAL UNIVERSITY OF SOUTH, C., MEMORIAL SLOAN KETTERING CANCER, C., MONTEFIORE MEDICAL, C., NANTOMICS, NATIONAL CANCER, I., NATIONAL HOSPITAL, A. N., NATIONAL HUMAN GENOME RESEARCH, I., NATIONAL INSTITUTE OF ENVIRONMENTAL HEALTH, S., NATIONAL INSTITUTE ON, D., OTHER COMMUNICATION, D., ONTARIO TUMOUR BANK, L. H. S. C., ONTARIO TUMOUR BANK, O. I. F. C. R., ONTARIO TUMOUR BANK, T. O. H., OREGON, H., SCIENCE, U., SAMUEL OSCHIN COMPREHENSIVE CANCER INSTITUTE, C.-S. M. C., INTERNATIONAL, S. R. A., ST JOSEPH’S CANDLER HEALTH, S., ELI, EDYTHE, L. B. I. O. M. I. O. T., HARVARD, U., RESEARCH INSTITUTE AT NATIONWIDE CHILDREN’S, H., SIDNEY KIMMEL COMPREHENSIVE CANCER CENTER AT JOHNS HOPKINS, U., UNIVERSITY OF, B., UNIVERSITY OF TEXAS, M. D. A. C. C., UNIVERSITY OF ABUJA TEACHING, H., UNIVERSITY OF ALABAMA AT, B., UNIVERSITY OF CALIFORNIA, I., UNIVERSITY OF CALIFORNIA SANTA, C., UNIVERSITY OF KANSAS MEDICAL, C., UNIVERSITY OF, L., UNIVERSITY OF NEW MEXICO HEALTH SCIENCES, C., UNIVERSITY OF NORTH CAROLINA AT CHAPEL, H., UNIVERSITY OF OKLAHOMA HEALTH SCIENCES, C., UNIVERSITY OF, P., UNIVERSITY OF SAO PAULO, R. A. P. M. S., UNIVERSITY OF SOUTHERN, C., UNIVERSITY OF, W., UNIVERSITY OF WISCONSIN SCHOOL OF, M., PUBLIC, H., VAN ANDEL RESEARCH, I. & WASHINGTON UNIVERSITY IN ST, L. 2017. Integrated genomic and molecular characterization of cervical cancer. Nature, 543, 378–384.

9. CANCER GENOME ATLAS RESEARCH, N., Weinstein, J. N., Collisson, E. A., Mills, G. B., Shaw, K. R., Ozenberger, B. A., Ellrott, K., Shmulevich, I., Sander, C. & Stuart, J. M. 2013. The Cancer Genome Atlas Pan-Cancer analysis project. Nat Genet, 45, 1113–20.

10. Carrara, M., Beccuti, M., Lazzarato, F., Cavallo, F., Cordero, F., Donatelli, S. & Calogero, R. A. 2013. State-of-the-art fusion-finder algorithms sensitivity and specificity. Biomed Res Int, 2013, 340620.

11. Chase, A., Ernst, T., Fiebig, A., Collins, A., Grand, F., Erben, P., Reiter, A., Schreiber, S. & Cross, N. C. 2010. Tfg, a target of chromosome translocations in lymphoma and soft tissue tumors, fuses to GPR128 in healthy individuals. Haematologica, 95, 20–6.

12. Clark, J., Rocques, P. J., Crew, A. J., Gill, S., Shipley, J., Chan, A. M., Gusterson, B. A. & Cooper, C. S. 1994. Identification of novel genes, SYT and Ssx, involved in the t(X;18)(p11.2;q11.2) translocation found in human synovial sarcoma. Nat Genet, 7, 502–8.

13. Collins, R. L., Brand, H., Karczewski, K. J., Zhao, X., Alfoldi, J., Francioli, L. C., Khera, A. V., Lowther, C., Gauthier, L. D., Wang, H., Watts, N. A., Solomonson, M., O’donnell-Luria, A., Baumann, A., Munshi, R., Walker, M., Whelan, C. W., Huang, Y., Brookings, T., Sharpe, T., Stone, M. R., Valkanas, E., Fu, J., Tiao, G., Laricchia, K. M., Ruano-Rubio, V., Stevens, C., Gupta, N., Cusick, C., Margolin, L., GENOME AGGREGATION DATABASE PRODUCTION, T., GENOME AGGREGATION DATABASE, C., Taylor, K. D., Lin, H. J., Rich, S. S., Post, W. S., Chen, Y. I., Rotter, J. I., Nusbaum, C., Philippakis, A., Lander, E., Gabriel, S., Neale, B. M., Kathiresan, S., Daly, M. J., Banks, E., Macarthur, D. G. & Talkowski, M. E. 2020. A structural variation reference for medical and population genetics. Nature, 581, 444–451.

14. COMMONS”, N. C. I. G. D. 2021. TCGA Study Abbreviations [Online]. Available: https://gdc.cancer.gov/resources-tcga-users/tcga-code-tables/tcga-study-abbreviations [Accessed].

15. CONSORTIUM, G. T. 2013. The Genotype-Tissue Expression (GTEx) project. Nat Genet, 45, 580–5.

16. Davidsen, J., Larsen, S., Coskun, M., Gogenur, I., Dahlgaard, K., Bennett, E. P. & Troelsen, J. T. 2018. The VTI1A-TCF4 colon cancer fusion protein is a dominant negative regulator of Wnt signaling and is transcriptionally regulated by intestinal homeodomain factor CDX2. PLoS One, 13, e0200215.

17. Dehghannasiri, R., Freeman, D. E., Jordanski, M., Hsieh, G. L., Damljanovic, A., Lehnert, E. & Salzman, J. 2019. Improved detection of gene fusions by applying statistical methods reveals oncogenic RNA cancer drivers. Proc Natl Acad Sci U S A, 116, 15524–15533.

18. Dobin, A., Davis, C. A., Schlesinger, F., Drenkow, J., Zaleski, C., Jha, S., Batut, P., Chaisson, M. & Gingeras, T. R. 2013. STAR: ultrafast universal RNA-seq aligner. Bioinformatics, 29, 15–21.

19. Edgren, H., Murumagi, A., Kangaspeska, S., Nicorici, D., Hongisto, V., Kleivi, K., Rye, I. H., Nyberg, S., Wolf, M., Borresen-Dale, A. L. & Kallioniemi, O. 2011. Identification of fusion genes in breast cancer by paired-end RNA-sequencing. Genome Biol, 12, R6.

20. Forbes, S. A., Beare, D., Boutselakis, H., Bamford, S., Bindal, N., Tate, J., Cole, C. G., Ward, S., Dawson, E., Ponting, L., Stefancsik, R., Harsha, B., Kok, C. Y., Jia, M., Jubb, H., Sondka, Z., Thompson, S., De, T. & Campbell, P. J. 2017. COSMIC: somatic cancer genetics at high-resolution. Nucleic Acids Res, 45, D777–D783.

21. Frattini, V., Trifonov, V., Chan, J. M., Castano, A., Lia, M., Abate, F., Keir, S. T., Ji, A. X., Zoppoli, P., Niola, F., Danussi, C., Dolgalev, I., Porrati, P., Pellegatta, S., Heguy, A., Gupta, G., Pisapia, D. J., Canoll, P., Bruce, J. N., Mclendon, R. E., Yan, H., Aldape, K., Finocchiaro, G., Mikkelsen, T., Prive, G. G., Bigner, D. D., Lasorella, A., Rabadan, R. & Iavarone, A. 2013. The integrated landscape of driver genomic alterations in glioblastoma. Nat Genet, 45, 1141–9.

22. Garalde, D. R., Snell, E. A., Jachimowicz, D., Sipos, B., Lloyd, J. H., Bruce, M., Pantic, N., Admassu, T., James, P., Warland, A., Jordan, M., Ciccone, J., Serra, S., Keenan, J., Martin, S., Mcneill, L., Wallace, E. J., Jayasinghe, L., Wright, C., Blasco, J., Young, S., Brocklebank, D., Juul, S., Clarke, J., Heron, A. J. & Turner, D. J. 2018. Highly parallel direct RNA sequencing on an array of nanopores. Nat Methods, 15, 201–206.

23. Gazendam, A. M., Popovic, S., Munir, S., Parasu, N., Wilson, D. & Ghert, M. 2021. Synovial Sarcoma: A Clinical Review. Curr Oncol, 28, 1909–1920.

24. Gingeras, T. R. 2009. Implications of chimaeric non-co-linear transcripts. Nature, 461, 206–11.

25. Goldman, M. J., Craft, B., Hastie, M., Repecka, K., Mcdade, F., Kamath, A., Banerjee, A., Luo, Y., Rogers, D., Brooks, A. N., Zhu, J. & Haussler, D. 2020. Visualizing and interpreting cancer genomics data via the Xena platform. Nat Biotechnol, 38, 675–678.

26. Grabherr, M. G., Haas, B. J., Yassour, M., Levin, J. Z., Thompson, D. A., Amit, I., Adiconis, X., Fan, L., Raychowdhury, R., Zeng, Q., Chen, Z., Mauceli, E., Hacohen, N., Gnirke, A., Rhind, N., DI Palma, F., Birren, B. W., Nusbaum, C., Lindblad-Toh, K., Friedman, N. & Regev, A. 2011. Full-length transcriptome assembly from RNA-Seq data without a reference genome. Nat Biotechnol, 29, 644–52.

27. Haas, B. 2021. Analyses, Code, and Data Supporting the FusionInspector Paper [Online]. Available: https://github.com/broadinstitute/FusionInspectorPaper [Accessed].

28. Haas, B., Dobin, A., Stransky, N., Li, B., Yang, X., Tickle, T., Bankapur, A., Ganote, C., Doak, T., Pochet, N., Sun, J., Wu, C., Gingeras, T. & Regev, A. 2017. STAR-Fusion: Fast and Accurate Fusion Transcript Detection from RNA-Seq. bioRxiv.

29. Haas, B. J. 2019. Trinity Cancer Transcriptome Analysis Toolkit [Online]. Available: https://github.com/NCIP/Trinity_CTAT/wiki [Accessed].

30. Haas, B. J., Dobin, A., Li, B., Stransky, N., Pochet, N. & Regev, A. 2019. Accuracy assessment of fusion transcript detection via read-mapping and de novo fusion transcript assembly-based methods. Genome Biology, 20, 213.

31. Haas, B. J., Papanicolaou, A., Yassour, M., Grabherr, M., Blood, P. D., Bowden, J., Couger, M. B., Eccles, D., Li, B., Lieber, M., Macmanes, M. D., Ott, M., Orvis, J., Pochet, N., Strozzi, F., Weeks, N., Westerman, R., William, T., Dewey, C. N., Henschel, R., Leduc, R. D., Friedman, N. & Regev, A. 2013. De novo transcript sequence reconstruction from RNA-seq using the Trinity platform for reference generation and analysis. Nat Protoc, 8, 1494–512.

32. Hale, R., Sandakly, S., Shipley, J. & Walters, Z. 2019. Epigenetic Targets in Synovial Sarcoma: A Mini-Review. Front Oncol, 9, 1078.

33. Hu, X., Wang, Q., Tang, M., Barthel, F., Amin, S., Yoshihara, K., Lang, F. M., Martinez-Ledesma, E., Lee, S. H., Zheng, S. & Verhaak, R. G. W. 2018. TumorFusions: an integrative resource for cancer-associated transcript fusions. Nucleic Acids Res, 46, D1144–D1149.

34. Jaiswal, S. & Ebert, B. L. 2019. Clonal hematopoiesis in human aging and disease. Science, 366.

35. Jerby-Arnon, L., Neftel, C., Shore, M. E., Weisman, H. R., Mathewson, N. D., Mcbride, M. J., Haas, B., Izar, B., Volorio, A., Boulay, G., Cironi, L., Richman, A. R., Broye, L. C., Gurski, J. M., Luo, C. C., Mylvaganam, R., Nguyen, L., Mei, S., Melms, J. C., Georgescu, C., Cohen, O., Buendia-Buendia, J. E., Segerstolpe, A., Sud, M., Cuoco, M. S., Labes, D., Gritsch, S., Zollinger, D. R., Ortogero, N., Beechem, J. M., Petur Nielsen, G., Chebib, I., Nguyen-Ngoc, T., Montemurro, M., Cote, G. M., Choy, E., Letovanec, I., Cherix, S., Wagle, N., Sorger, P. K., Haynes, A. B., Mullen, J. T., Stamenkovic, I., Rivera, M. N., Kadoch, C., Wucherpfennig, K. W., Rozenblatt-Rosen, O., Suva, M. L., Riggi, N. & Regev, A. 2021. Opposing immune and genetic mechanisms shape oncogenic programs in synovial sarcoma. Nat Med, 27, 289–300.

36. Jin, K., Wang, S., Zhang, Y., Xia, M., Mo, Y., Li, X., Li, G., Zeng, Z., Xiong, W. & He, Y. 2019. Long non-coding RNA PVT1 interacts with MYC and its downstream molecules to synergistically promote tumorigenesis. Cell Mol Life Sci, 76, 4275–4289.

37. Kasif, S. & Roberts, R. J. 2020. We need to keep a reproducible trace of facts, predictions, and hypotheses from gene to function in the era of big data. PLoS Biol, 18, e3000999.

38. Kim, P., Yiya, K. & Zhou, X. 2020. FGviewer: an online visualization tool for functional features of human fusion genes. Nucleic Acids Res, 48, W313–W320.

39. Kim, P., Yoon, S., Kim, N., Lee, S., Ko, M., Lee, H., Kang, H., Kim, J. & Lee, S. 2010. ChimerDB 2.0--a knowledgebase for fusion genes updated. Nucleic Acids Res, 38, D81–5.

40. Klijn, C., Durinck, S., Stawiski, E. W., Haverty, P. M., Jiang, Z., Liu, H., Degenhardt, J., Mayba, O., Gnad, F., Liu, J., Pau, G., Reeder, J., Cao, Y., Mukhyala, K., Selvaraj, S. K., Yu, M., Zynda, G. J., Brauer, M. J., Wu, T. D., Gentleman, R. C., Manning, G., Yauch, R. L., Bourgon, R., Stokoe, D., Modrusan, Z., Neve, R. M., DE Sauvage, F. J., Settleman, J., Seshagiri, S. & Zhang, Z. 2015. A comprehensive transcriptional portrait of human cancer cell lines. Nat Biotechnol, 33, 306–12.

41. Kumar, S., Razzaq, S. K., Vo, A. D., Gautam, M. & Li, H. 2016a. Identifying fusion transcripts using next generation sequencing. Wiley Interdiscip Rev Rna, 7, 811–823.

42. Kumar, S., Vo, A. D., Qin, F. & Li, H. 2016b. Comparative assessment of methods for the fusion transcripts detection from RNA-Seq data. Sci Rep, 6, 21597.

43. Kurzrock, R., Gutterman, J. U. & Talpaz, M. 1988. The molecular genetics of Philadelphia chromosome-positive leukemias. N Engl J Med, 319, 990–8.

44. Lagstad, S., Zhao, S., Hoff, A. M., Johannessen, B., Lingjaerde, O. C. & Skotheim, R. I. 2017. chimeraviz: a tool for visualizing chimeric RNA. Bioinformatics, 33, 2954–2956.

45. Lasorella, A., Sanson, M. & Iavarone, A. 2017. FGFR-TACC gene fusions in human glioma. Neuro Oncol, 19, 475–483.

46. Liquori, A., Ibanez, M., Sargas, C., Sanz, M. A., Barragan, E. & Cervera, J. 2020. Acute Promyelocytic Leukemia: A Constellation of Molecular Events around a Single PML-RARA Fusion Gene. Cancers (Basel), 12.

47. Liu, Q., Hu, Y., Stucky, A., Fang, L., Zhong, J. F. & Wang, K. 2020. LongGF: computational algorithm and software tool for fast and accurate detection of gene fusions by long-read transcriptome sequencing. BMC Genomics, 21, 793.

48. Neuendorff, N. R., Burmeister, T., Dorken, B. & Westermann, J. 2016. BCR-ABL-positive acute myeloid leukemia: a new entity? Analysis of clinical and molecular features. Ann Hematol, 95, 1211–21.

49. Nome, T., Hoff, A. M., Bakken, A. C., Rognum, T. O., Nesbakken, A. & Skotheim, R. I. 2014. High frequency of fusion transcripts involving TCF7L2 in colorectal cancer: novel fusion partner and splice variants. PLoS One, 9, e91264.

50. Northcott, P. A., Shih, D. J., Peacock, J., Garzia, L., Morrissy, A. S., Zichner, T., Stutz, A. M., Korshunov, A., Reimand, J., Schumacher, S. E., Beroukhim, R., Ellison, D. W., Marshall, C. R., Lionel, A. C., Mack, S., Dubuc, A., Yao, Y., Ramaswamy, V., Luu, B., Rolider, A., Cavalli, F. M., Wang, X., Remke, M., Wu, X., Chiu, R. Y., Chu, A., Chuah, E., Corbett, R. D., Hoad, G. R., Jackman, S. D., Li, Y., Lo, A., Mungall, K. L., Nip, K. M., Qian, J. Q., Raymond, A. G., Thiessen, N. T., Varhol, R. J., Birol, I., Moore, R. A., Mungall, A. J., Holt, R., Kawauchi, D., Roussel, M. F., Kool, M., Jones, D. T., Witt, H., Fernandez, L. A., Kenney, A. M., Wechsler-Reya, R. J., Dirks, P., Aviv, T., Grajkowska, W. A., Perek-Polnik, M., Haberler, C. C., Delattre, O., Reynaud, S. S., Doz, F. F., Pernet-Fattet, S. S., Cho, B. K., Kim, S. K., Wang, K. C., Scheurlen, W., Eberhart, C. G., Fevre-Montange, M., Jouvet, A., Pollack, I. F., Fan, X., Muraszko, K. M., Gillespie, G. Y., Di Rocco, C., Massimi, L., Michiels, E. M., Kloosterhof, N. K., French, P. J., Kros, J. M., Olson, J. M., Ellenbogen, R. G., Zitterbart, K., Kren, L., Thompson, R. C., Cooper, M. K., Lach, B., Mclendon, R. E., Bigner, D. D., Fontebasso, A., Albrecht, S., Jabado, N., Lindsey, J. C., Bailey, S., Gupta, N., Weiss, W. A., Bognar, L., Klekner, A., VAN Meter, T. E., Kumabe, T., Tominaga, T., Elbabaa, S. K., Leonard, J. R., Rubin, J. B., et al. 2012. Subgroup-specific structural variation across 1,000 medulloblastoma genomes. Nature, 488, 49–56.

51. Peng, Z., Yuan, C., Zellmer, L., Liu, S., Xu, N. & Liao, D. J. 2015. Hypothesis: Artifacts, Including Spurious Chimeric RNAs with a Short Homologous Sequence, Caused by Consecutive Reverse Transcriptions and Endogenous Random Primers. J Cancer, 6, 555–67.

52. Rautiainen, M., Durai, D. A., Chen, Y., Xin, L., Low, H. M., Göke, J., Marschall, T. & Schulz, M. H. 2020. AERON: Transcript quantification and gene-fusion detection using long reads. bioRxiv, 2020.01.27.921338.

53. Ren, R. 2005. Mechanisms of BCR-ABL in the pathogenesis of chronic myelogenous leukaemia. Nat Rev Cancer, 5, 172–83.

54. Robinson, J. 2019. igv-reports [Online]. Available: https://github.com/igvteam/igv-reports [Accessed].

55. Rubin, M. A., Maher, C. A. & Chinnaiyan, A. M. 2011. Common gene rearrangements in prostate cancer. J Clin Oncol, 29, 3659–68.

56. Sabir, S. R., Yeoh, S., Jackson, G. & Bayliss, R. 2017. EML4-ALK Variants: Biological and Molecular Properties, and the Implications for Patients. Cancers (Basel), 9.

57. Schmidt, B., Cmero, M., Ekert, P., Davidson, N. & Oshlack, A. 2021. Slinker: Visualising novel splicing events in RNA-Seq data. F1000Res, 10, 1255.

58. Schmidt, B. M., Davidson, N. M., Hawkins, A. D. K., Bartolo, R., Majewski, I. J., Ekert, P. G. & Oshlack, A. 2018. Clinker: visualizing fusion genes detected in RNA-seq data. Gigascience, 7.

59. Shivram, H. & Iyer, V. R. 2018. Identification and removal of sequencing artifacts produced by mispriming during reverse transcription in multiple RNA-seq technologies. Rna, 24, 1266–1274.

60. Soda, M., Choi, Y. L., Enomoto, M., Takada, S., Yamashita, Y., Ishikawa, S., Fujiwara, S., Watanabe, H., Kurashina, K., Hatanaka, H., Bando, M., Ohno, S., Ishikawa, Y., Aburatani, H., Niki, T., Sohara, Y., Sugiyama, Y. & Mano, H. 2007. Identification of the transforming EML4-ALK fusion gene in non-small-cell lung cancer. Nature, 448, 561–6.

61. TERRA. Terra: a scalable platform for biomedical research [Online]. Available: https://terra.bio/ [Accessed].

62. Tolomeo, D., Agostini, A., Visci, G., Traversa, D. & Storlazzi, C. T. 2021. PVT1: A long non-coding RNA recurrently involved in neoplasia-associated fusion transcripts. Gene, 779, 145497.

63. Tomlins, S. A., Rhodes, D. R., Perner, S., Dhanasekaran, S. M., Mehra, R., Sun, X. W., Varambally, S., Cao, X., Tchinda, J., Kuefer, R., Lee, C., Montie, J. E., Shah, R. B., Pienta, K. J., Rubin, M. A. & Chinnaiyan, A. M. 2005. Recurrent fusion of TMPRSS2 and ETS transcription factor genes in prostate cancer. Science, 310, 644–8.

64. Traag, V. A., Waltman, L. & Van Eck, N. J. 2019. From Louvain to Leiden: guaranteeing well-connected communities. Sci Rep, 9, 5233.

65. Wang, Q., Xia, J., Jia, P., Pao, W. & Zhao, Z. 2013. Application of next generation sequencing to human gene fusion detection: computational tools, features and perspectives. Brief Bioinform, 14, 506–19.

66. Wei, Z., Zhou, C., Zhang, Z., Guan, M., Zhang, C., Liu, Z. & Liu, Q. 2019. The Landscape of Tumor Fusion Neoantigens: A Pan-Cancer Analysis. iScience, 21, 249–260.

67. Wright, M. N. & Ziegler, A. 2017. ranger: A Fast Implementation of Random Forests for High Dimensional Data in C++ and R. 2017 %9 C++; classification; machine learning; R; random forests; Rcpp; recursive partitioning; survival analysis %! ranger: A Fast Implementation of Random Forests for High Dimensional Data in C++ and R, 77, 17.

68. Yan, M., Wang, J., Ren, Y., Li, L., He, W., Zhang, Y., Liu, T. & Li, Z. 2019. Over-expression of FSIP1 promotes breast cancer progression and confers resistance to docetaxel via MRP1 stabilization. Cell Death Dis, 10, 204.

69. Yang, W., Lee, K. W., Srivastava, R. M., Kuo, F., Krishna, C., Chowell, D., Makarov, V., Hoen, D., Dalin, M. G., Wexler, L., Ghossein, R., Katabi, N., Nadeem, Z., Cohen, M. A., Tian, S. K., Robine, N., Arora, K., Geiger, H., Agius, P., Bouvier, N., Huberman, K., Vanness, K., Havel, J. J., Sims, J. S., Samstein, R. M., Mandal, R., Tepe, J., Ganly, I., Ho, A. L., Riaz, N., Wong, R. J., Shukla, N., Chan, T. A. & Morris, L. G. T. 2019. Immunogenic neoantigens derived from gene fusions stimulate T cell responses. Nat Med, 25, 767–775.

70. Yoshihara, K., Wang, Q., Torres-Garcia, W., Zheng, S., Vegesna, R., Kim, H. & Verhaak, R. G. 2015. The landscape and therapeutic relevance of cancer-associated transcript fusions. Oncogene, 34, 4845–54.

71. Yu, C. Y., Liu, H. J., Hung, L. Y., Kuo, H. C. & Chuang, T. J. 2014. Is an observed non-co-linear RNA product spliced in trans, in cis or just in vitro? Nucleic Acids Res, 42, 9410–23.

72. Zhang, J., Gao, T. & Maher, C. A. 2017. INTEGRATE-Vis: a tool for comprehensive gene fusion visualization. Sci Rep, 7, 17808.

73. Zhou, J. X., Yang, X., Ning, S., Wang, L., Wang, K., Zhang, Y., Yuan, F., Li, F., Zhuo, D. D., Tang, L. & Zhuo, D. 2017. Identification of KANSARL as the first cancer predisposition fusion gene specific to the population of European ancestry origin. Oncotarget, 8, 50594–50607.

74. Zhou, L., Liu, R., Liang, X., Zhang, S., Bi, W., Yang, M., He, Y., Jin, J., Li, S., Yang, X., Fu, J. & Zhang, P. 2020. lncRNA RP11-624L4.1 Is Associated with Unfavorable Prognosis and Promotes Proliferation via the CDK4/6-Cyclin D1-Rb-E2F1 Pathway in NPC. Mol Ther Nucleic Acids, 22, 1025–1039.

