## Supplemental Figures for "Targeted *in silico* characterization of fusion transcripts in tumor and normal tissues via FusionInspector"

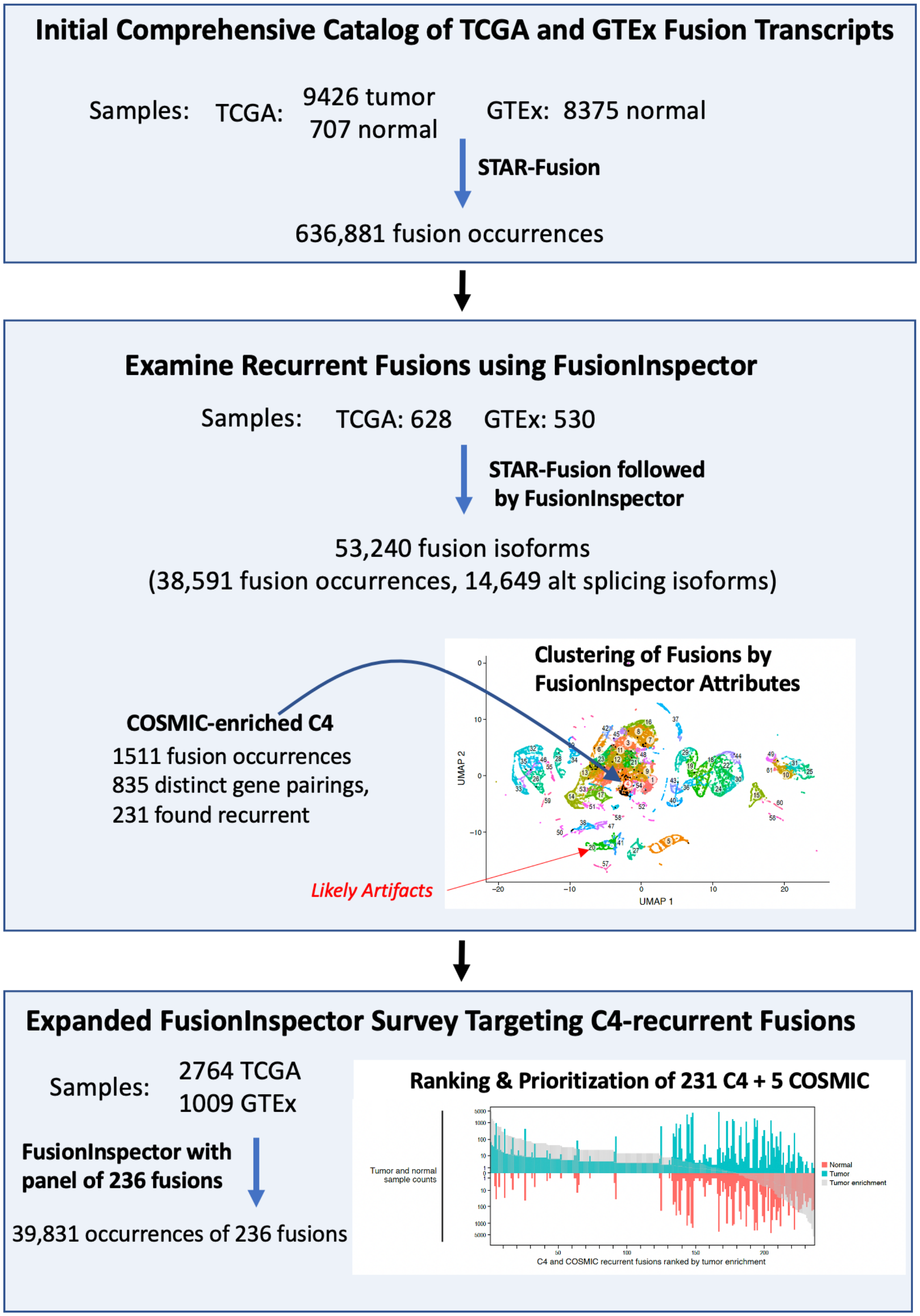


**Figure S1. Roadmap for our Intertwined Development and Application of FusionInspector.**

From an initial comprehensive catalog of fusion transcripts predicted from TCGA and GTEx (**top**), we selected a subset of samples representative of recurrent fusions, which we further investigated using FusionInspector (**center**). Clustering of inspected fusion transcript isoforms based on FusionInspector computed attributes coupled with annotation of known cancer fusions yielded our discovery of COSMIC-fusion enriched cluster C4. Certain other clusters had features consistent with likely artifacts. Using fusion cluster assignments, we built a random forest classifier to predict new fusion instances according to cluster labels based on their FusionInspector attributes, ultimately yielding predictions of new fusion isoforms as COSMIC-like, artifact-like, or other fusion cluster type. To gain further insights into additional COSMIC-like fusions that might be prioritized for further study, we targeted FusionInspector to numerous additional predicted instances of these C4 fusions across TCGA and GTEx samples (**bottom**). We examined predicted fusion type classifications as predominantly COSMIC-like, artifact-like, or other, coupled with functional impacts on coding sequences, prevalence of occurrence across normal and tumor samples, and we prioritized fusions accordingly, demonstrating the utility of FusionInspector for in silico characterization and prioritization of predicted fusion transcripts while identifying novel fusion isoforms of potential relevance to tumor or normal biology.

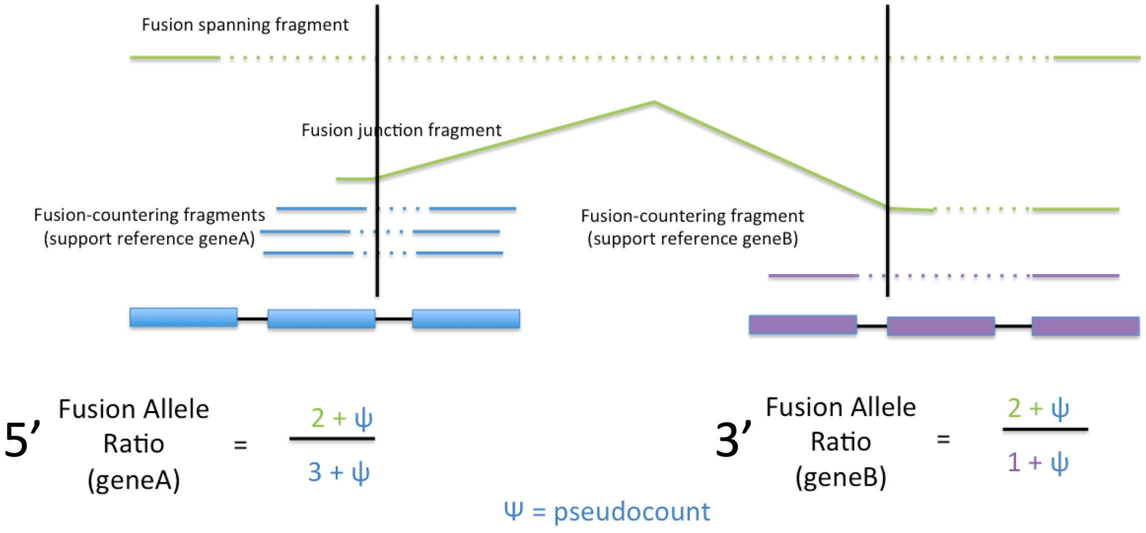


**Figure S2. Computing 5' and 3' fusion allelic ratios (FAR):** RNA-seq reads supporting the fusion (green) are compared to the RNA-seq reads supporting the unfused partner transcripts (5': blue; 3': red) at the breakpoint to compute 5'- and 3'-gene fusion allelic ratios. In practice, we use pseudocount=1.

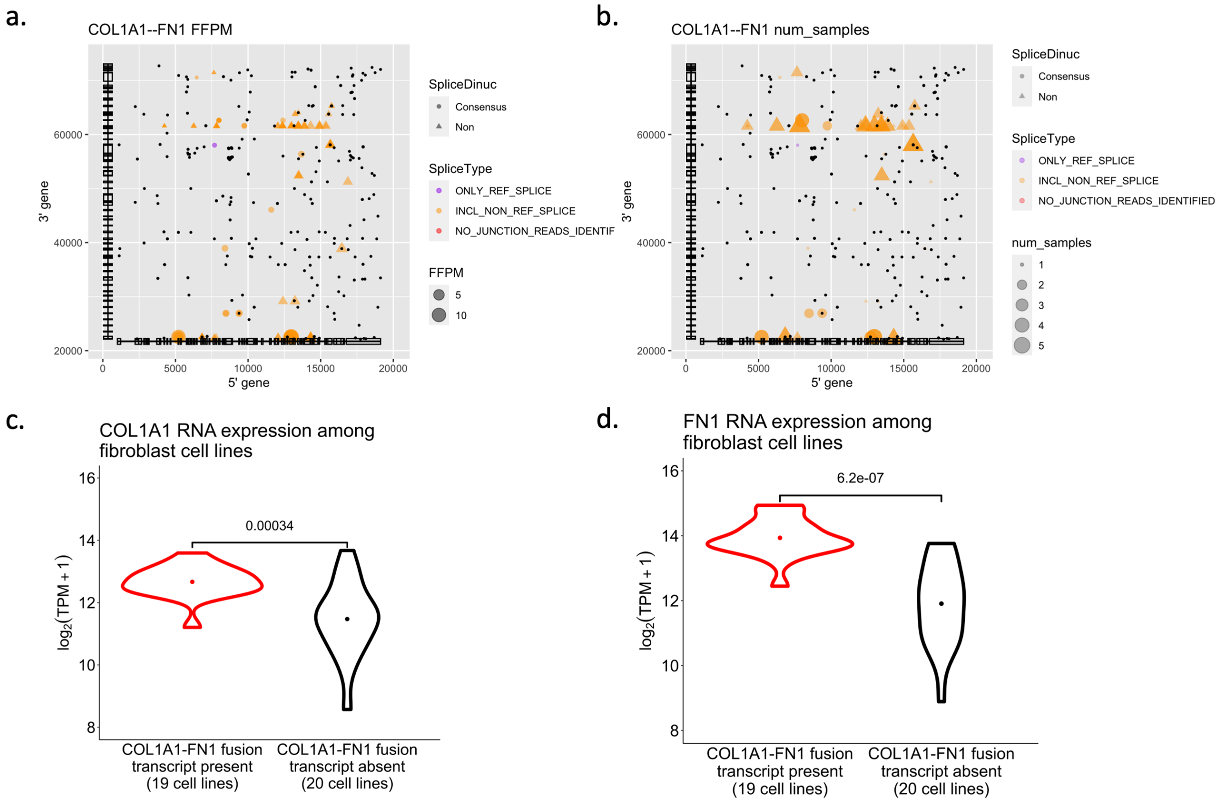


**Figure S3: Fusion transcript COL1A1--FN1 identified recurrently among cancer associated fibroblast cell lines in the Cancer Cell Line Encyclopedia is likely artifactual.** (**a,b**) Isoform expression levels for putative fusions (**a,** dot size) or number of samples (**b**, dot size), splice type (dot color) and splice junction dinucleotide (dot shape) at each fusion breakpoint position involving the 5’ (x axis) and 3’ (y axis) partners of COL1A1—FN1 in five fibroblast cell lines with highest COL1A1--FN1 fusion read support (HS600T_FIBROBLAST, HS688AT_FIBROBLAST, HS739T_FIBROBLAST, HS819T_FIBROBLAST, and HS822T_FIBROBLAST). Black dots: positions of microhomology (10 base exact match). Structures of collapsed isoforms for fusion partner genes are drawn along each axis. (**c,d**) Distribution of (c) COL1A1 or (d) FN1 expression levels (x axis) in fibroblast cell lines with (red) or without (black) a detected COL1A1--FN1 fusion. Expression data are derived from DepMap (https://depmap.org/portal/download/) release 19q2. Because most of the reads contributing to the robust FFPM are spanning fragments, most breakpoints occur at non-canonical dinucleotide splice sites, and there is significant microhomology detected at these breakpoints, COL1A1-FN1 is likely an RT-PCR artifact resulting from increased expression of partner genes, as opposed to the molecular lesion.

a.

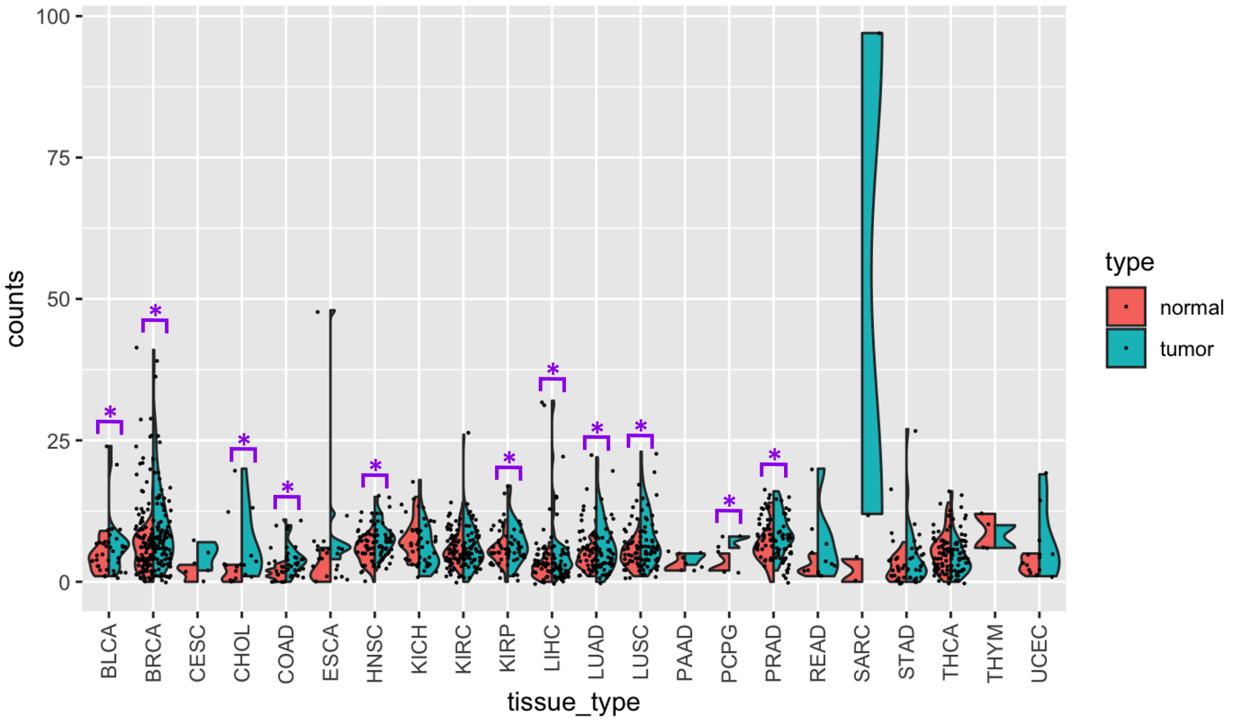


b.

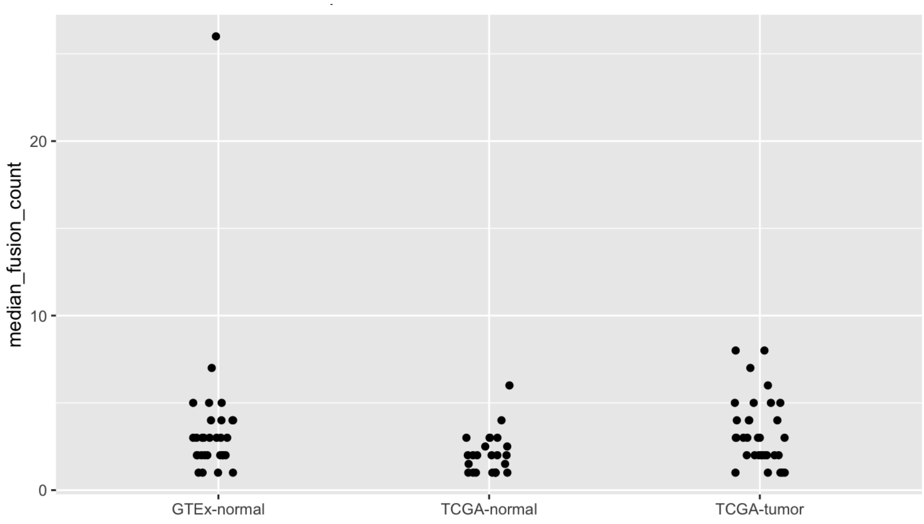


**Figure S4: Higher number of fusion predictions in tumor vs. matched normal samples in TCGA but not GTEx**. (**a**) Higher number of fusions in tumors vs. matched normal samples in multiple tumor types in TCGA. Distribution of number of fusions (y axis) in each tumor (blue) and matched normal (red) tissue type (x axis) from TCGA, for those fusions where there is at least one fusion in either tumor or matched normal sample at 0.1 FFPM threshold. * Benjamini-Hochberg FDR < 0.05, one-sided paired t-test. (**b**) Similar number (p-value=0.6, t-test) of leniently predicted fusions in TCGA and GTEx. Median number of fusions (y axis) predicted per tumor or normal tissue type (dots) (x axis) after filtering for a minimum of 0.1 FFPM and removing likely readthrough transcripts involving co-linear neighboring genes within 100kb. with typically four to five fusions predicted per normal or tumor tissue type. In normal tissues, pancreas is an outlier (median of 27 fusions per sample). Fusions were identified with lenient evidence requirements to allow sensitive detection: a minimum of only two supporting reads with at least one defining the fusion breakpoint, and disabling annotation-based filters (**Methods**).

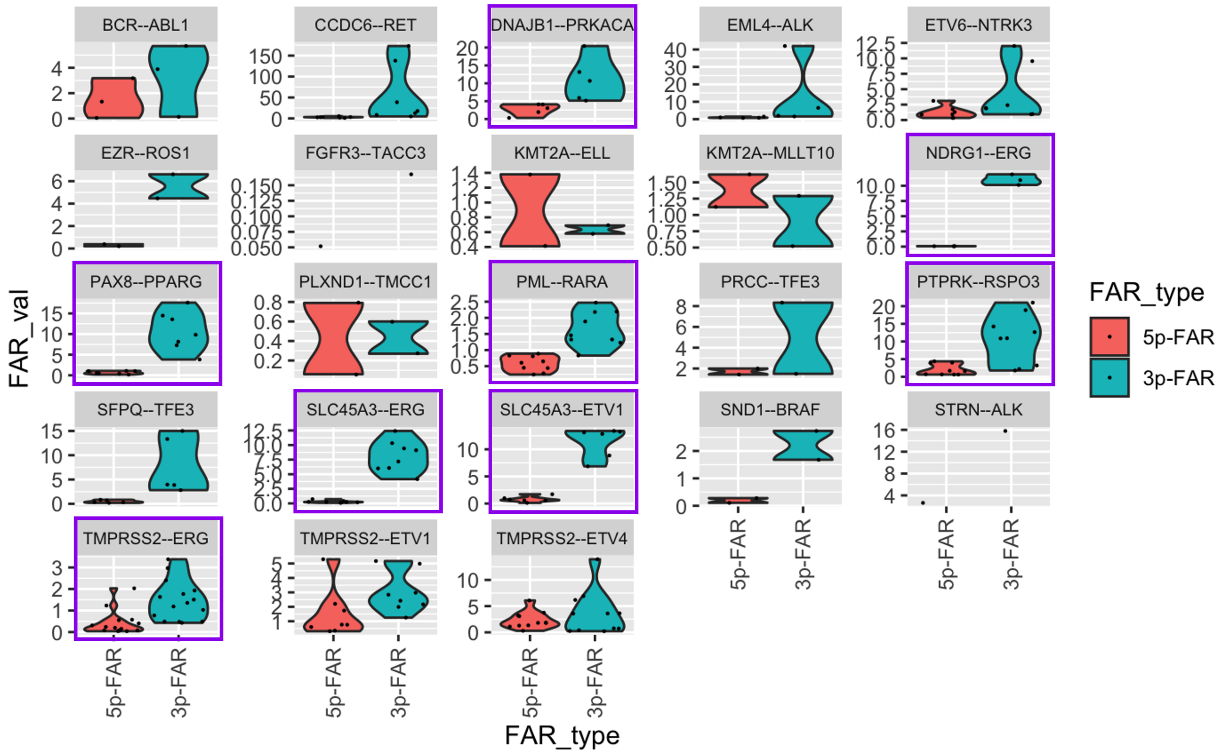


**Figure S5. Low 5’ vs. 3’ FAR in COSMIC fusions in C4.**

Distribution of fusion allele ratio (FAR, y axis) for the 5’ (red) and 3’ (blue) FAR for 23 representative COSMIC fusions in Cluster C4. Purple: Benjamini-Hochberg FDR < 0.05, t test. Exceptions (*e.g.*, with KMT2A as the 5' partner gene) have generally low FAR values for both partner genes.

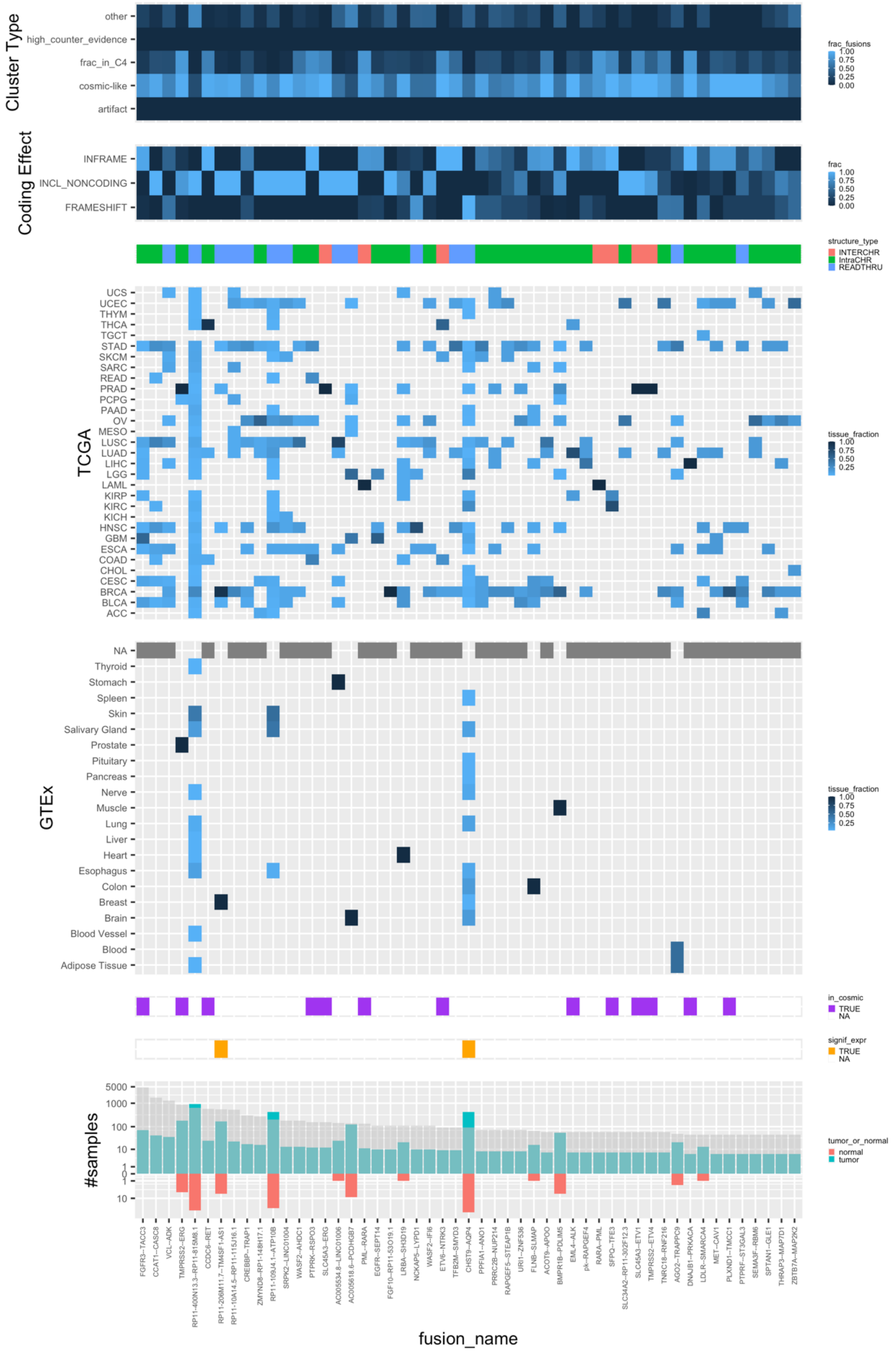


**Figure S6a (fusions 1-50)**

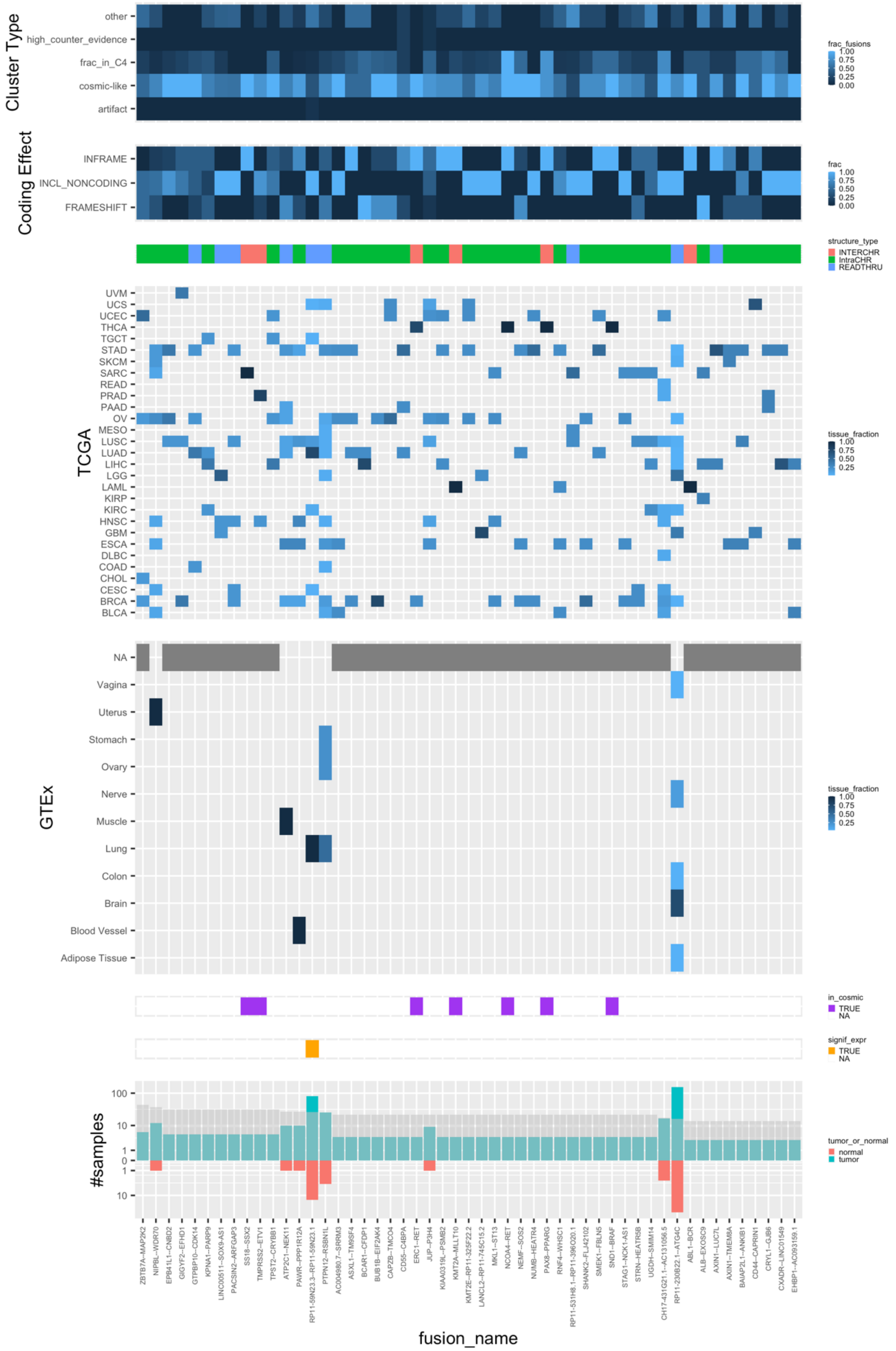


**Figure S6b (fusions 51-100)**

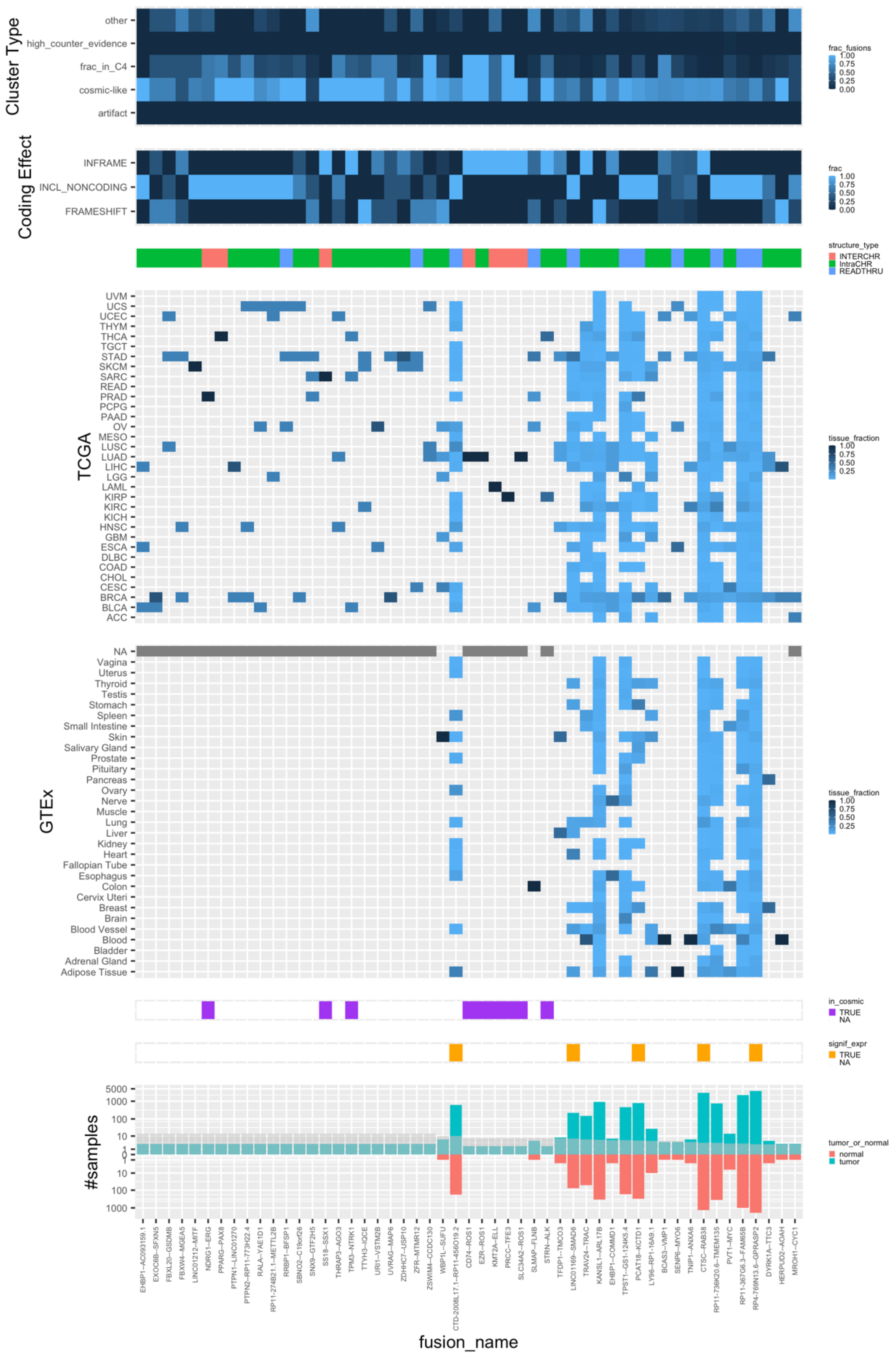


**Figure S6c (fusions: 101-150)**

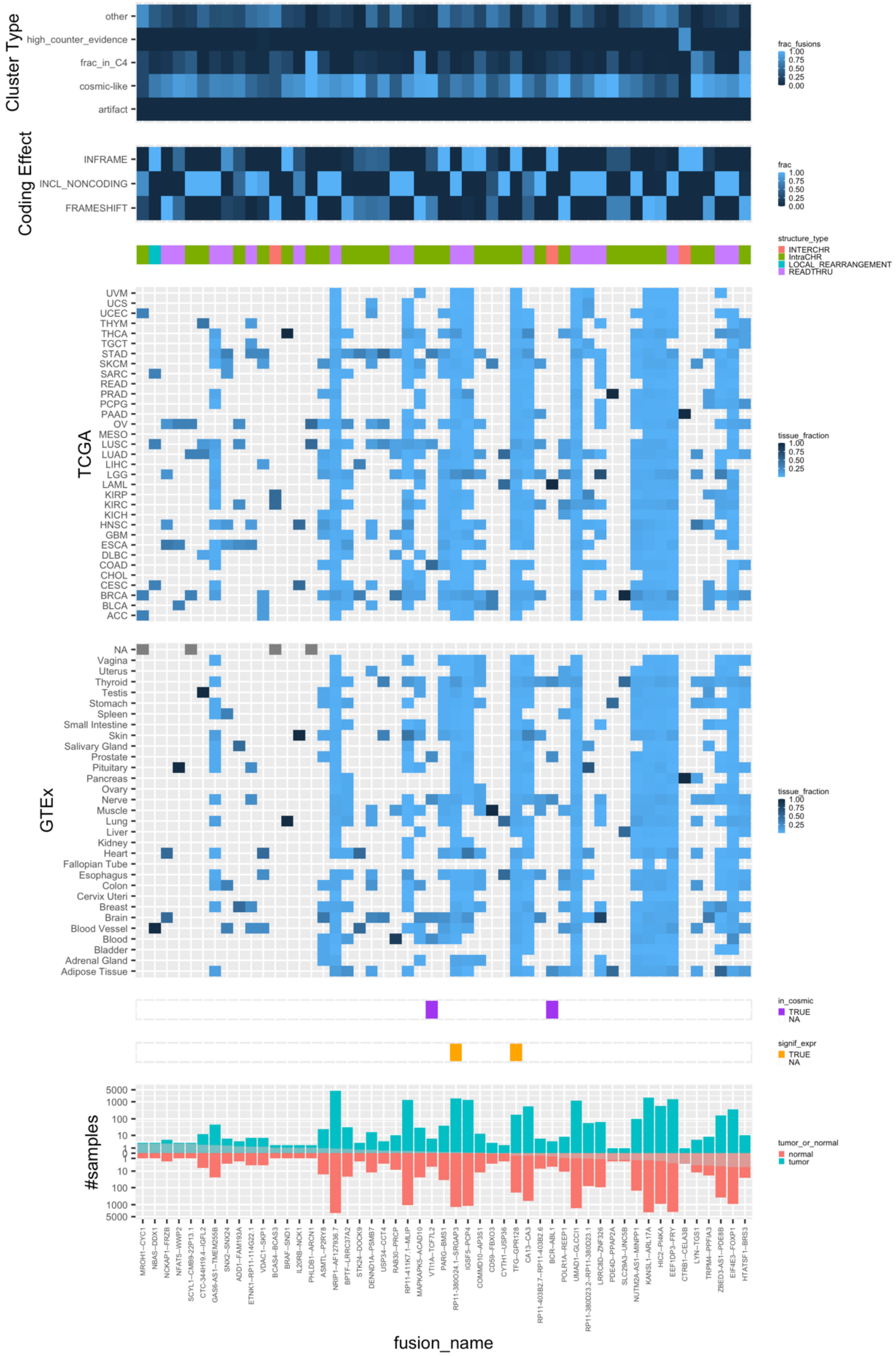


**Figure S6d (fusions: 151-200)**

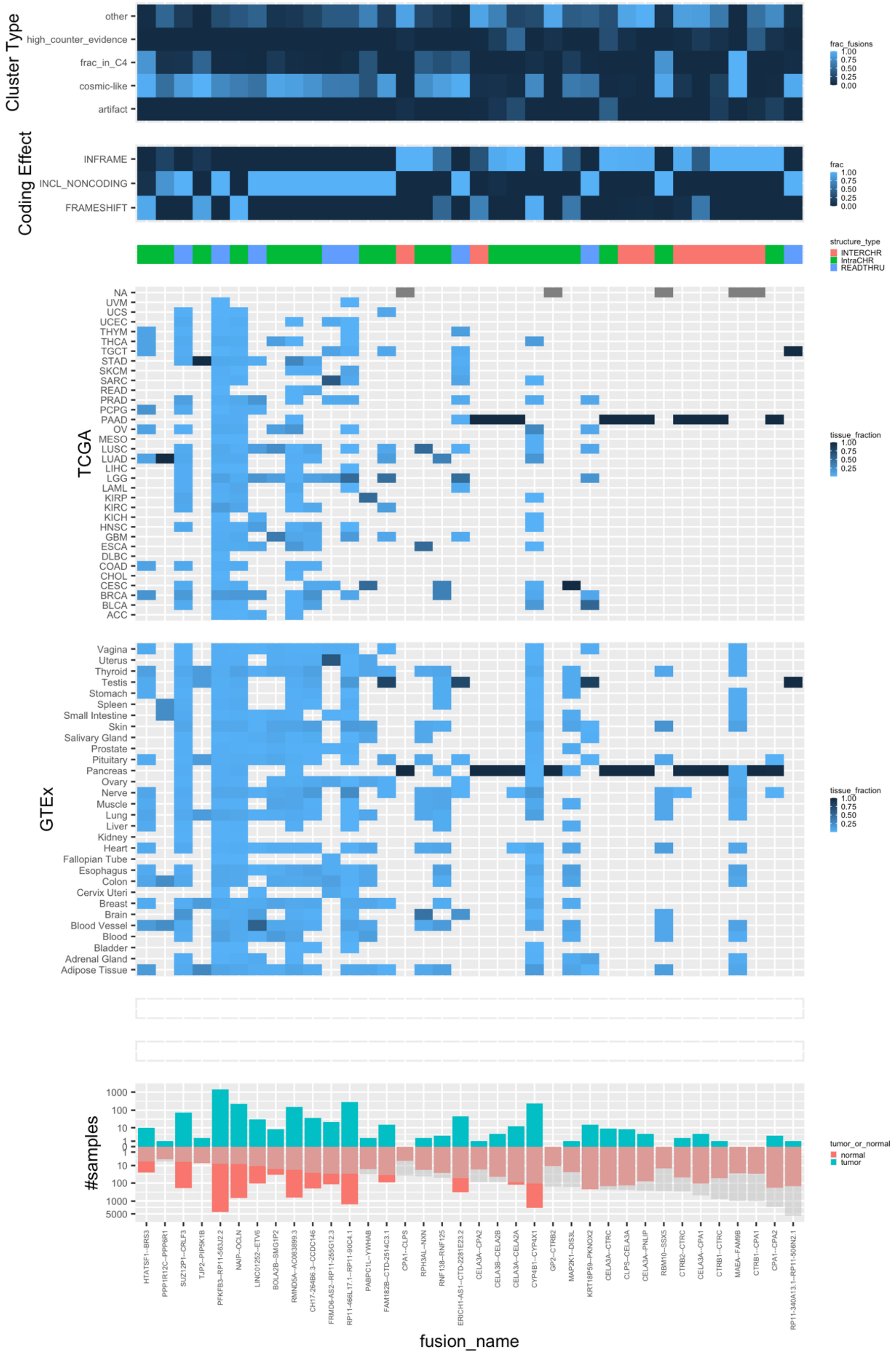


**Figure S6e (fusions: 201-236)**

**Figure S6. Properties of recurrent C4 and COSMIC fusions, ranked by tumor/normal sample occurrence ratio.** 236 selected COSMIC-peak-enriched (C4) and additional COSMIC fusions (columns / x axis), rank ordered by tumor enrichment and shown in ~50 fusion increments (**a-e**) with fraction of the instances of each fusion in each category based on predicted Leiden cluster labels (top panel 3 rows) or corresponding to presumed impact on coding sequence (top panel, bottom rows); fusion structure type based on the fusion partner's chromosomal location (second from top); fraction of instances that is in each tumor or tissue type in TCGA and GTEx (third from top, rows); presence in COSMIC (third from bottom, purple), significantly higher expression in tumors *vs*. normal tissues (second fro bottom**,** Wilcoxon rank sum test applied to FFPM, Benjamini Hochberg FDR $< 0.05$ and median tumor FFPM > median normal FFPM, orange), number of tumor (seagreen) or normal (light red) samples (bottom**,** y axis) predicted by STAR-Fusion to contain the fusion, rank ordered by tumor enrichment (bottom, x axis, **Methods**, gray).

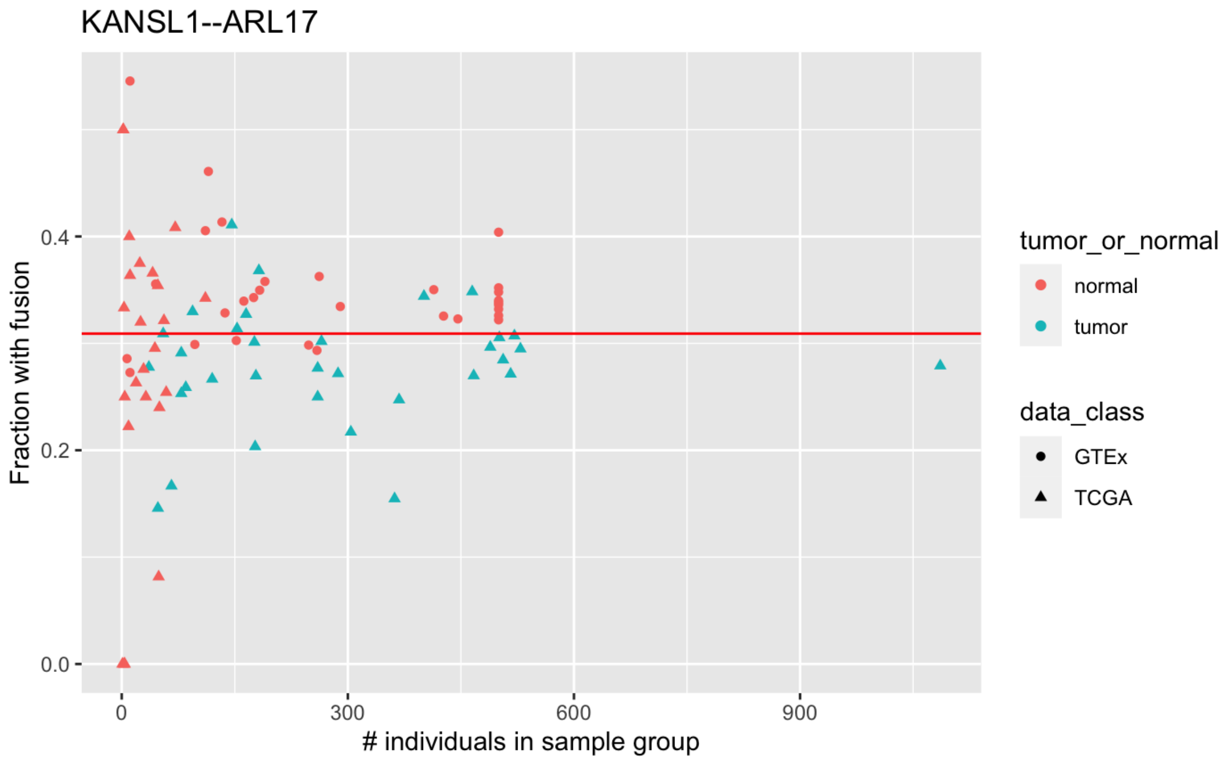


**Figure S7: Fraction of TCGA and GTEx tissue samples containing evidence of fusion transcript KANSL1--ARL17.** Each data point corresponds to the fraction of corresponding tissue samples demonstrating evidence of the KANSL1--ARL17 fusion (called as KANSL1--ARL17A or KANSL1--ARL17B). No minimum FFPM threshold applied.

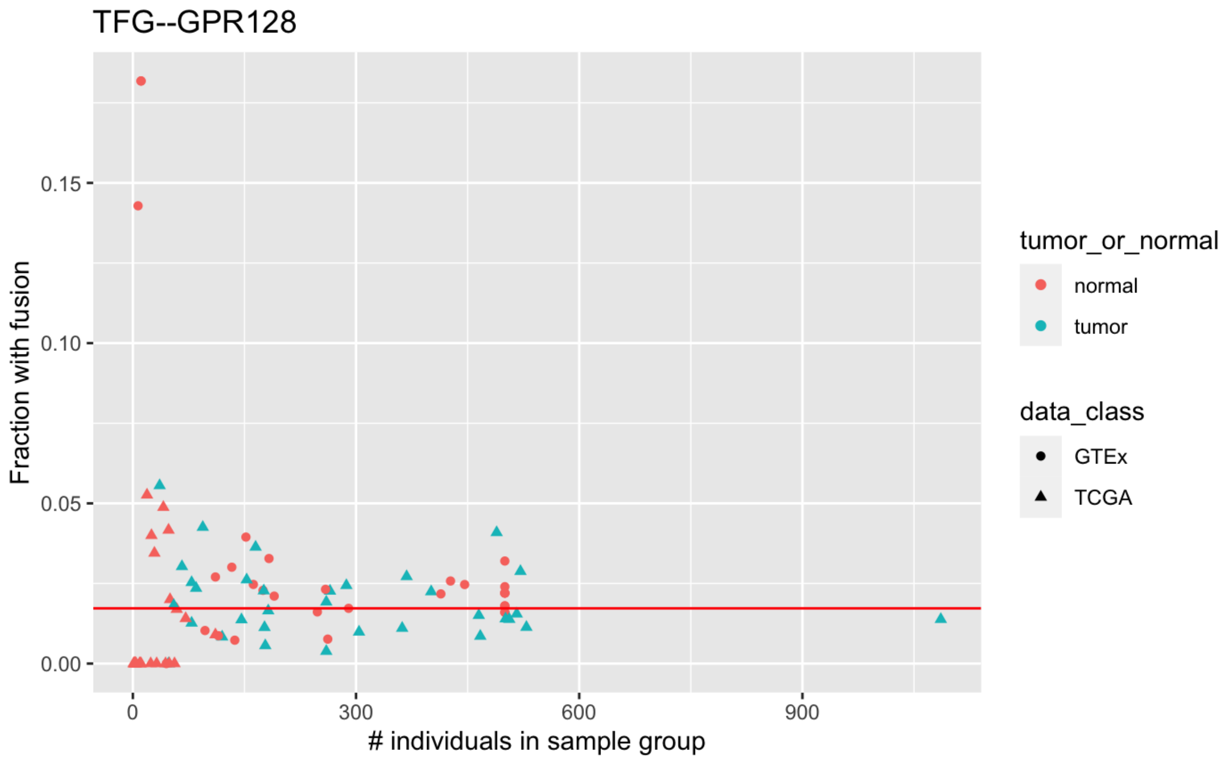


**Figure S8: Evidence of fusion transcript TFG--GPR128 across TCGA and GTex**. Fraction of tumor (blue) or normal tissue (red) samples (y axis) with evidence of the TFG--GPR128 fusion transcript, associated with a known germline structural variant estimated with a European population allele frequency of ~2%. No minimum FFPM threshold applied.

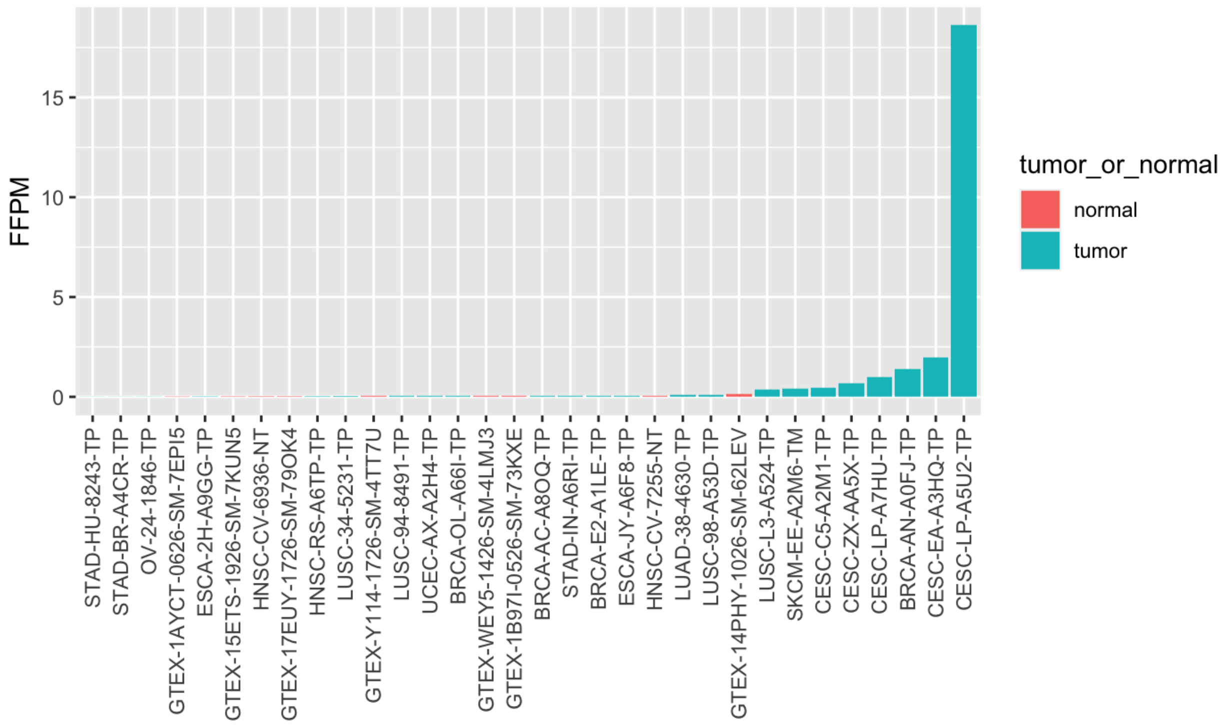


**Figure S9. Higher PVT1--MYC fusion expression in tumors vs. normal samples, especially in cervical cancer samples.**

Expression level (FFPM) of PVT1--MYC fusion in each sample (x axis) in which it is detected. Only the top 11/32 samples have FusionInspector estimated FFPM>0.1, five of which correspond to cervical cancer (CESC), and corresponds to a hotspot site of HPV insertion in cervical cancer genomes.

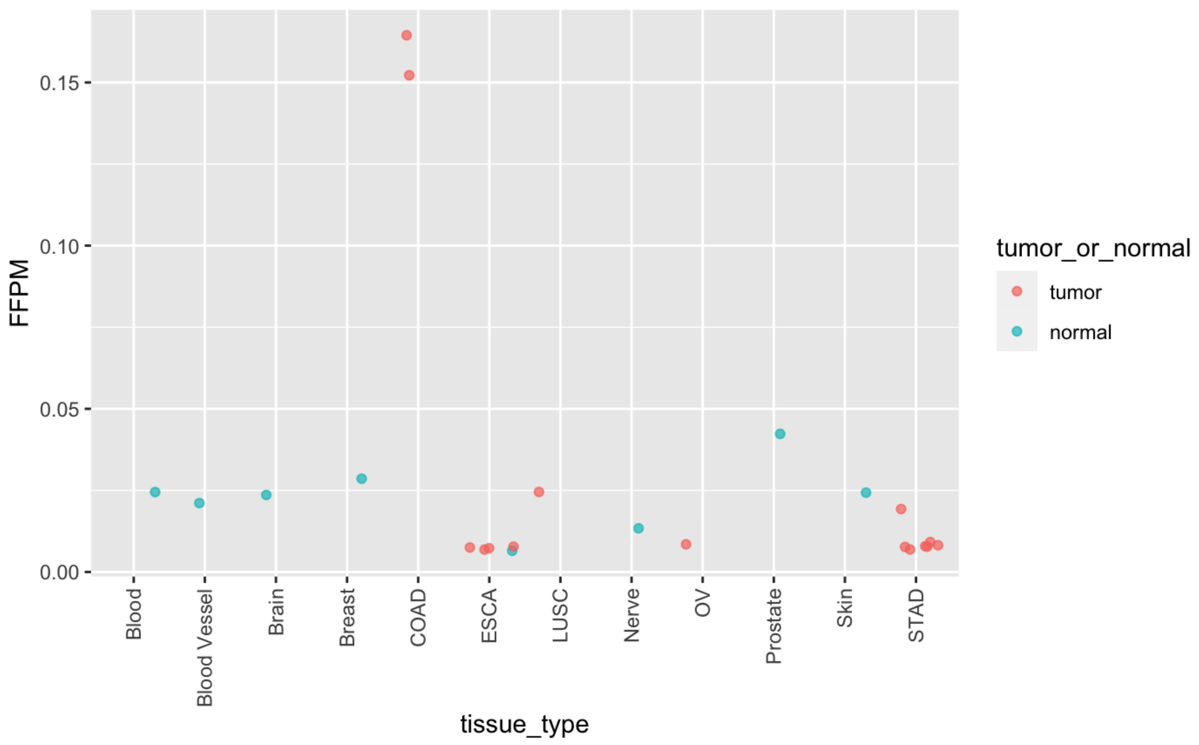


**Figure S10. The VTI1A--TCF7L2 fusion is highly expressed in colon cancer.**

Expression level (FFPM) of VTI1A--TCF7L2 fusion in each normal (blue) or tumor (red) sample (*x* axis) in which it is detected (no FFPM threshold applied), ordered by sample type.

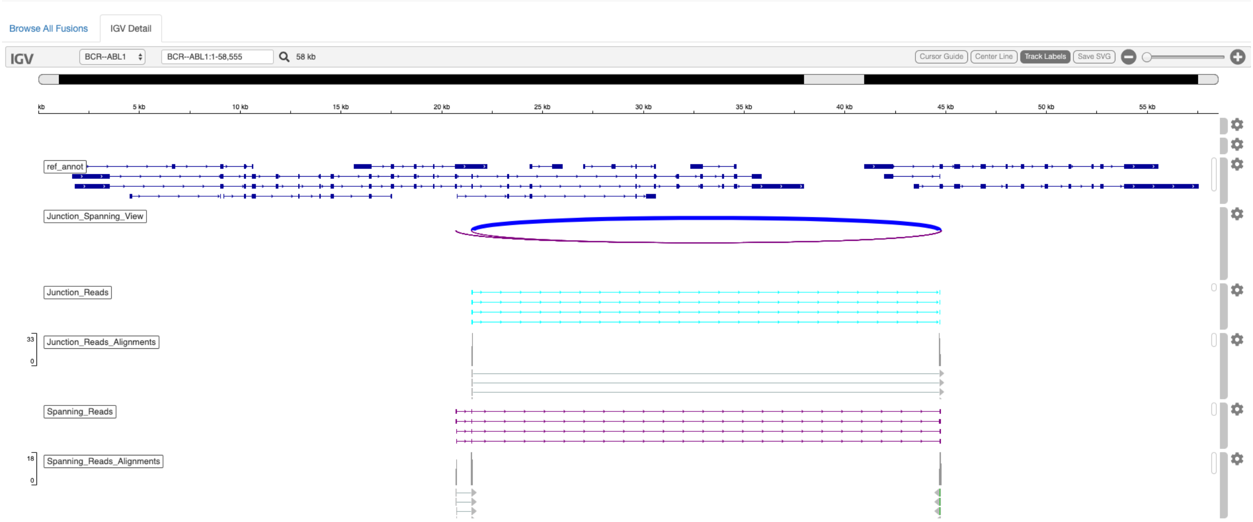


**Figure S11: FusionInspector visualization leveraging igv-reports**.

Fusion evidence reads are shown for fusion BCR--ABL1 aligned to the FusionInspector fusion contig in comparison to the reference gene structures. The FusionInspector igv-reports are stand-alone html application files provided as and output file and convenient for rapid interactive data exploration in any computing framework including cloud-computing architectures such as Terra, or popular platforms such as Galaxy.

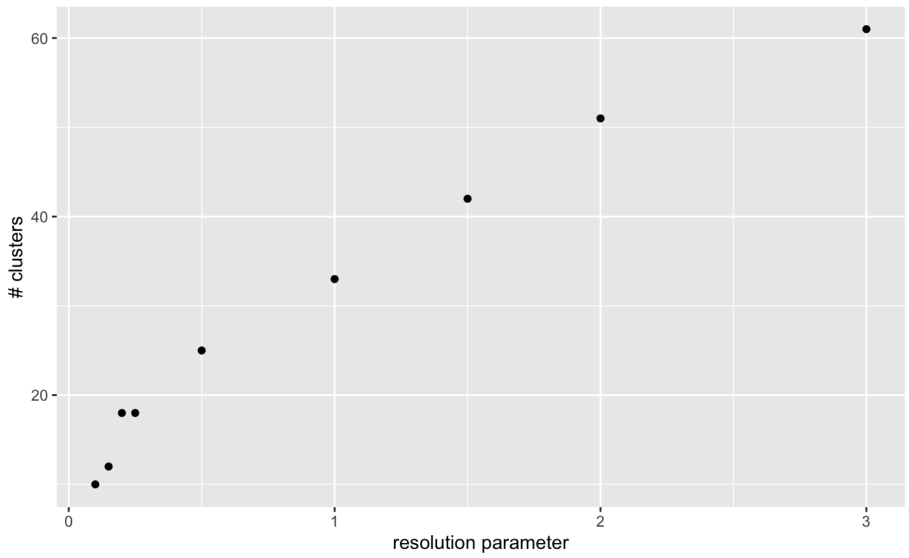


a.

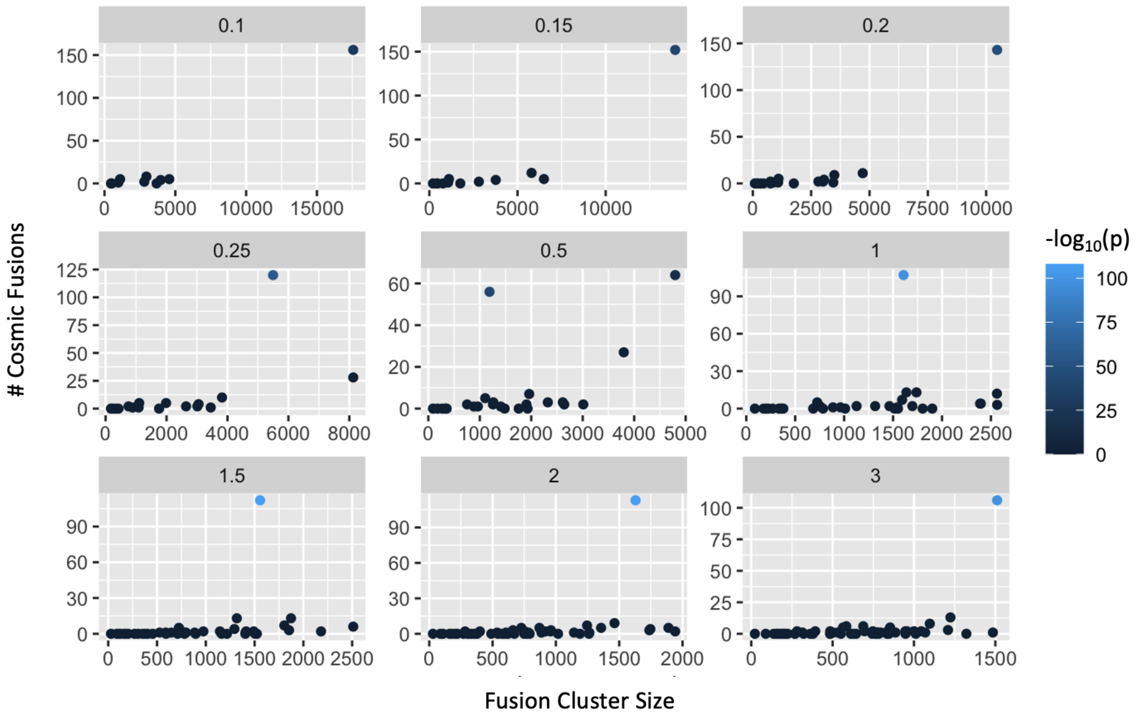


b.

**Figure S12. Number and sizes of fusion clusters *vs*. Leiden resolution setting.**

(**a**) Increasing number of clusters at higher resolution parameters. Number of clusters (*y* axis) at each Leiden resolution parameter value (*x* axis). (**b**) Cluster size (x axis) and the number of

COSMIC fusions (y axis) that are members of the cluster for each cluster (dot). Color: $-\log_{10}$(P-value) of enrichment for COSMIC fusion membership (Fisher's exact test, one-sided).
